# An improved plant toolset for high-throughput recombineering

**DOI:** 10.1101/659276

**Authors:** J. Brumos, C. Zhao, Y. Gong, D. Soriano, A.P. Patel, M.A. Perez-Amador, A.N. Stepanova, J.M Alonso

## Abstract

Gene functional studies often rely on the expression of a gene of interest as transcriptional and translational fusions with specialized tags. Ideally, this is done in the native chromosomal contexts to avoid potential misexpression artifacts. Although recent improvements in genome editing make it possible to directly modify the target genes in their native chromosomal location, classical transgenesis is still the preferred experimental approach chosen in most gene tagging studies because of its time efficiency and accessibility. We have developed a recombineering-based tagging system that brings together the convenience of the classical transgenic approaches and the high degree of confidence in the obtained results provided by the direct chromosomal tagging achievable by genome editing strategies. These simple and customizable recombineering toolsets and protocols allow for high-throughput generation of a variety of genetic modifications. In addition, a highly efficient recombinase-mediated cassette exchange system has been developed to facilitate the transfer of the desired sequences from a BAC clone to a transformation-compatible binary vector, expanding the use of the recombineering approaches beyond *Arabidopsis*. The utility of this system is demonstrated by the generation of over 250 whole-gene translational fusions and 123 *Arabidopsis* transgenic lines corresponding to 62 auxin-related genes, and the characterization of the translational reporter expression patterns for 14 auxin biosynthesis genes.

## INTRODUCTION

The last few years have witnessed dramatic advances in high-throughput experimental and computational approaches to investigate the molecular mechanisms behind biological processes. Nevertheless, certain types of information-rich functional data are still exceedingly tedious and time-consuming to obtain. Thus, any experimental approaches that require *in vivo* expression of the gene of interest (GOI) to, for example, gather high-resolution spatiotemporal expression patterns, determine protein subcellular localization, or identify protein-protein and protein-DNA/RNA complexes, still heavily rely on classical restriction enzyme or recombination-based cloning strategies. Although these classical approaches are simple and accessible and, therefore, widely used, they have several limitations regarding scalability and may suffer from uncertainty when trying to capture the native expression patterns and levels of the genes under investigation. This uncertainty comes from the need to choose the DNA sequences to be included in the construct with the risk that some unknown, but important, regulatory sequences may be left out. This is not a trivial problem when the native expression pattern of a GOI needs to be imposed on the tagged gene. In the absence of a strict criterium, more or less arbitrary lengths of DNA sequences (typically from 1 to 4 kb of sequences upstream and 1 kb or less of sequences downstream of the start and stop codon, respectively) or all of the intergenic sequences flanking the GOI, are usually chosen. These strategies, however, do not guarantee that all regulatory sequences are captured. Genetic complementation of a mutant line is relied upon to support that the expression patterns of the generated transgene accurately reflect that of the corresponding native gene. This time consuming and not fully foolproof approach is, however, not possible when either a mutant line is not available or, what is most common, when the mutant does not display any detectable phenotype. The obvious solution to this problem is to increase the size of the sequences flanking the GOI that would be included in the transgene or, even better, to insert the tag or the desired modification in the GOI directly in its native chromosomal location. Although the latter genome-editing approach is highly desirable and the number of reports of precise gene editing in plants is constantly increasing ((Cermak et al., 2015; Begemann et al., 2017; Yu et al., 2017; Dahan-Meir et al., 2018; Li et al., 2018) and reviewed in (Soyars et al., 2018), the transgenic approach is still the most widely used methodology to generate plants expressing genes carrying a tag or other modifications that facilitate their visualization or biochemical characterization. Classical transgenic approaches are not ideal either, as they become tedious and inefficient as the size of the DNA fragments used increases.

To overcome the limitations of traditional transgenic approaches, highly efficient homologous recombination in bacterial strains engineered to express the Exo, Beta and Gam proteins from the lambda phage, also known as lambda red recombineering system (Yu et al., 2000; Copeland et al., 2001), have been developed. The high efficiency of this recombineering system has made it an essential tool in bacterial genome engineering (Isaacs et al., 2011) allowing for the rapid, efficient and simultaneous editing of hundreds of loci in the bacterial genomes. Although the lambda red system has not been shown to work in eukaryotic cells, DNA from higher organisms can be efficiently modified using this system when introduced into recombineering-ready *E. coli* strains. Thus, recombineering has been successfully used to generate genome-wide collections of fluorescently tagged proteins in several model organisms, such as *Drosophila* and *C. elegans* (Sarov et al., 2012; Sarov et al., 2016). In addition to the *E. coli* recombineering strains (Warming et al., 2005), several other system-specific elements are required in order to make this technology accessible to a research community. First of all, a collection of sequence-indexed genomic clones covering the whole genome of the organism of interest needs to be available. This is essential to easily identify a clone containing a GOI and the flanking sequences containing all of the putative regulatory sequences for that gene. In the case of plants, the reintroduction of these large genomic DNA fragments into the plant genome typically requires the use of *Agrobacterium*-mediated transformation. This imposes an additional requirement that the vector carrying the large genomic DNA fragments should be compatible with *Agrobacterium*-mediated transformation. Alternatively, the large DNA fragments from a bacterial artificial chromosome (BAC) would need to be transferred to a suitable binary vector (Bitrian et al., 2011). In addition, unrestricted availability of a set of reusable recombineering cassettes suitable for the insertion of tags commonly used in plant research at any position in any GOI, as well as tools that allow for the generation of custom-designed tagging cassettes or the introduction of any other sequence modifications in the genes of interest, are essential for the popularization of this technology among plant biologists. Finally, robust and simple protocols to facilitate the use of recombineering in any plant biology research lab with a standard molecular biology setup, as well as scalable pipelines that allow for the implementation of this technology to entire gene families, pathways or the even the whole genome is essential for the plant community to take full advantage of the benefits offered by the recombineering technology.

Previously, we have shown that recombineering could be used to generate whole-gene translational fusions and point mutations in genes harbored in transformation-ready bacterial artificial chromosomes (TACs) and that these large TAC clones could be used for *Agrobacterium*-mediated transformation (Zhou et al., 2011). However, this original system has several limitations. First, it requires a sequence-indexed collection of TAC clones, in practice restricting its use to *Arabidopsis*. It also employs classical recombineering cassettes based on the selectable *galK* system (Warming et al., 2005) that relies on specialized media and expensive reagents (Warming et al., 2005). In addition, the relatively low efficiency of the contra-selection steps used to replace the *galK* gene by the tag of interest precludes this approach from being scaled up and requires significant troubleshooting when first adopted in a lab. Herein, we present a new set of tools and protocols that overcome all of these limitations. Namely, the new plant recombineering kit we describe here allows for the use of standard media and antibiotic selection, it provides a set of ready-to-use tags and a vector that can be utilized to convert any tag of interest into a recombineering-ready cassette. Importantly, a new set of plasmids and cassettes has been generated to facilitate the transfer of tens of thousands of base pairs from a BAC to a high-capacity binary vector, opening this technology to many plant species for which sequence-indexed genomic clones covering the genome are available. Finally, we have compiled sequence information from two *Arabidopsis* TAC libraries into a public genome browser allowing for the easy identification of TAC clones containing the *Arabidopsis* GOI. All of the vectors and cassettes required to carry out recombineering experiments in plants are available via the ABRC, while the JAtY and Kazusa TAC libraries (Hirose et al., 2015) are available from the ABRC and RIKEN BRC public stock centers. To demonstrate the utility of this system, we have tagged over 250 genes with different tags. We have made publicly available 123 transgenic lines corresponding to 62 genes through the ABRC and NASC. Among these lines are those corresponding to *GUS* translational fusions of all members of the *TAA1/TAR* and *YUC* auxin biosynthetic enzyme families implicated in the production of auxin, indole-3-acetic acid (IAA), from amino acid tryptophan via indole-3-pyruvic acid (IPyA). The characterization of these lines in the roots and hypocotyls of seedlings grown under different pharmacological treatments, as well as in untreated inflorescences and flowers, provides a detailed and comprehensive map of the auxin biosynthetic machinery.

## RESULTS

### Generation of excisable antibiotic-based recombineering cassettes

Classical recombineering strategies (Warming et al., 2005; Zhou et al., 2011) rely on two consecutive recombineering steps. In the first step, a positive-negative selectable marker such as *galK* is inserted in the genomic location to be modified, followed by a second recombineering step where *galK* is substituted by the desired tag or replacement sequence (Figure 1). One drawback of this time-consuming approach is that the negative selection step is prone to false positives (Warming et al., 2005; Zhou et al., 2011) and often several colonies per construct need to be tested to identify a true recombination product with the desired changes. An alternative approach to reduce the number of recombineering reactions needed has been the use of bifunctional recombineering cassettes that contain both the tag to be inserted in the GOI and a positive/negative selectable marker. This selectable marker is flanked by *flipase* (*FLP*) recognition target sites (*FRT*s) (Tursun et al., 2009) (Figure 1), thus enabling marker removal by activating the expression of a *FLP* recombinase with very high efficiency (Warming et al., 2005). In this alternative recombineering system, the positive selectable marker is first used to identify insertion events of the recombineering cassette in the GOI. An inducible *FLP* recombinase already engineered in some *E. coli* recombineering strains, such as SW105, is then used to trigger the excision of the selectable marker leaving behind just the reporter gene and a 36 nt *FRT* scar. The identification of these excision events can be facilitated by the loss of the *galK* activity that in the presence of 2-deoxy galactose inhibits bacterial growth (Warming et al., 2005). With the final goal to facilitate the use of recombineering and to allow for increased throughput, we have also adopted and improved the bifunctional cassettes containing both a selectable marker and a tag of interest. Although initially we chose the classical *galK* selectable marker to generate these bifunctional cassettes due to its counter-selectable capabilities (Warming et al., 2005; Zhou et al., 2011), we later generated a simplified and easier-to-use antibiotic-based excisable bifunctional recombineering cassette (Alonso and Stepanova, 2014) to better exploit the high efficiency of the *FLP*-based excision system already engineered into the *SW105* recombineering strain genome. Here, we have expanded the collection of bifunctional recombineering cassettes to a total of 11. These new antibiotic-based cassettes consist of several of the most commonly used tags in plants and an antibiotic resistance gene flanked by the *FRT* sites (Figure1, Table S5). In addition to simplifying and accelerating the selection of recombination events, these ampicillin- or tetracycline-based recombineering cassettes are compatible not only with the selectable markers of two end-sequenced TAC libraries in *Arabidopsis* (Zhou et al., 2011; Hirose et al., 2015), but also with the most popular plant BAC libraries that use kanamycin or chloramphenicol as the antibiotic selection in bacteria (Budiman et al., 2000; Yuan et al., 2000; Zhang et al., 2016). In addition, several fluorescent protein gene and *GUS* tags incorporated in these new cassettes have been codon-optimized for high expression in plants (Table S5).

**Figure 1.**
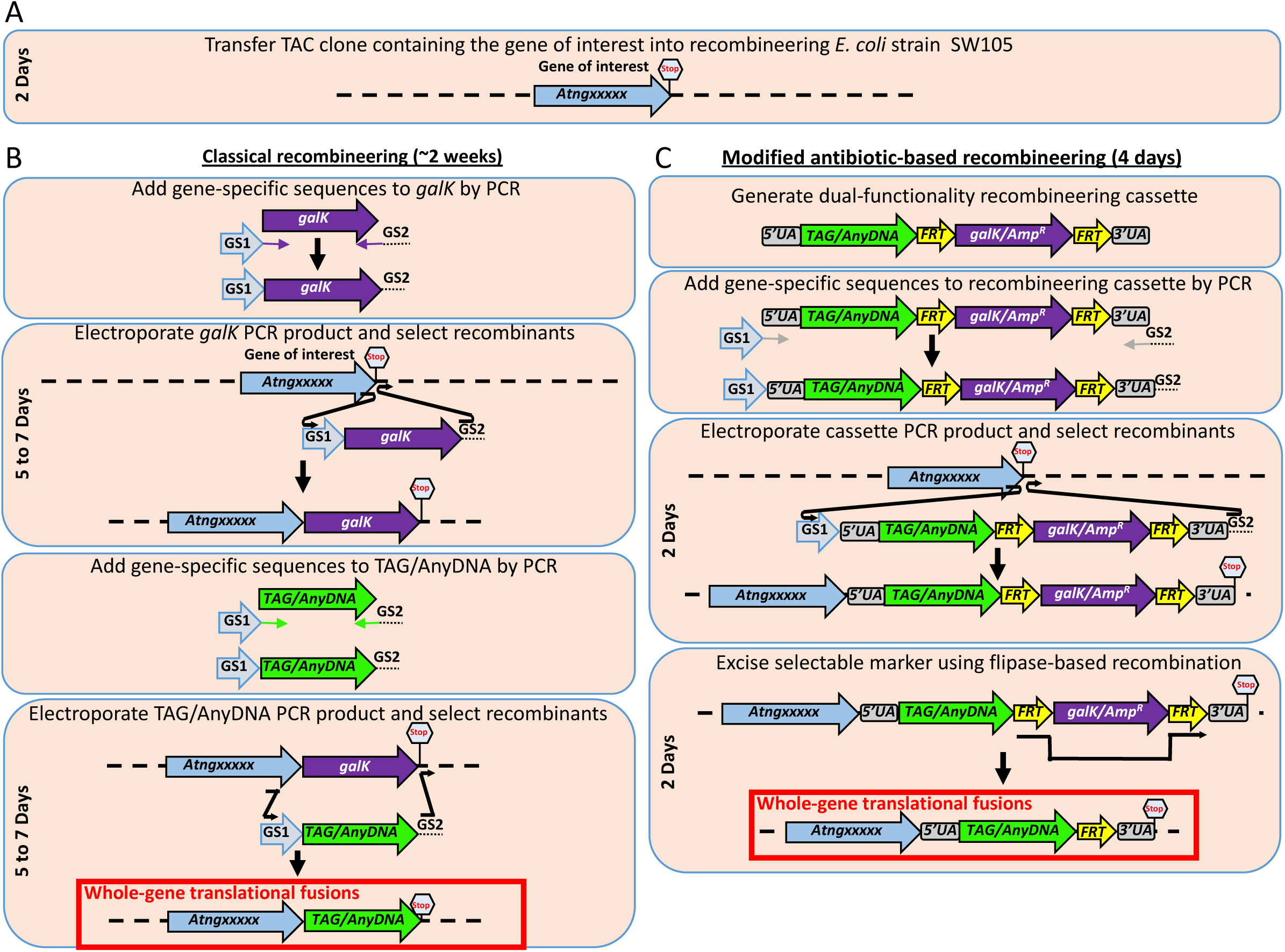
Timeline comparison between the classical and new accelerated recombineering. (A) The first step in any recombineering experiment is the identification of a genomic clone (typically a TAC or a BAC) containing the gene or sequences of interest. **(B)** In the classical *galK*-based system, the *galK* positive/negative selectable marker is amplified using a pair of primers that contain at least 40 nucleotides of sequence corresponding to the sequence flanking the desired insertion site in the target genomic DNA clone. In this example, the amplification of the *galK* cassette with the GS1 and GS2 primers will result in the production of an amplicon (*GS1-galK-GS2*) that will target the *galK* selectable marker to the 3’ of the gene just before the stop codon. The electroporation of this amplicon in a recombineering competent *E. coli* strain such as SW105 and the selection of the *galK-*positive colonies will result in a clone containing the *galK* marker just before the stop codon in the gene of interest (second panel). Using the same set of primers used to amplify the *galK* cassette, a *TAG/AnyDNA* cassette (such as *GFP*) is amplified (third panel) and used to replace *galK* by the *TAG/AnyDNA* sequence (bottom panel). This sequence replacement can be accomplished by electroporating the *GS1-TAG/AnyDNA-GS2* amplicon into the recombineering cells carrying the gene of interest tagged with *galK* and selecting for clones that lost *galK* in minimum media supplemented with 2-deoxy-galactose. Only *galK-*negative colonies will survive in the presence of this chemical. **(C)** The faster and user-friendly bifunctional cassette system combines the selectable marker (such as *galK* or an antibiotic resistance gene) and the tag of interest in a single cassette (top panel). By flanking the sequences of the selectable marker with the flipase (FLP) recognition target sites (*FRT*s), the selectable marker sequence can be readily removed post-insertion by a highly efficient *in vivo* FLP reaction. Similarly to the classical approach, the bifunctional large cassette, *GS1-5’UA-TAG/AnyDNA-FRT-galK/AmpR-FRT-5’UA-GS2* is first amplified with a pair of primers, GS1 and GS2 (second panel), to add the gene-specific sequences that will target the recombineering cassette to the desired location in the gene. By electroporating this cassette into the recombineering *E. coli* strain SW105 containing the gene of interest and selecting for, in this example, ampicillin-resistant clones, the bacteria with the desired construct can be efficiently and rapidly identified (third panel). Finally, the induction of FLP recombinase already engineered in the *SW105* strain would result in the removal of the sequences corresponding to the selectable marker (bottom panel), leading to the tag containing the reporter or epitope of interest followed by a 36-nt-long *FRT*-containing scar that encodes 12 extra amino acids. The approximate time period required in each step is indicated. The GS1 primer should have the following structure: 5’-40nt just upstream of the nucleotide after which you want to insert your tag followed by the *5’UA* sequence -ggaggtggaggtggagct -3’. Similarly, the GS2 primer should have the structure: 5’-40nt corresponding to the reverse complement of the sequence just downstream of the nucleotide in front of which you want to insert your tag followed by the 3’UA sequence ggccccagcggccgcagcagcacc-3’.

Importantly, all our recombineering cassettes share the same 5’ and 3’ universal adaptor sequences (Table S5). These sequences common to all our constructs serve two purposes. On the one hand, these adaptor sequences allow for the use of the same set of gene-specific 60mer primers to tag a GOI with any of the different tags in the collection. On the other hand, the in-frame adaptor sequences encode a poly-glycine and a poly-alanine linker, providing a flexible connection, and thus minimizing conformational interferences between the protein of interest and the corresponding tag (Tian et al., 2004). Finally, these adaptors have been designed to allow the same cassettes to be used in N-terminal, C-terminal or internal translational fusion experiments.

Although the new antibiotic-based recombineering cassettes make the generation of the translational fusions much simpler and more efficient, they do not allow for the same level of flexibility as provided by the classical *galK* system. Thus, for example, the counter-selectable properties of *galK* can be used, once inserted in the GOI, to generate replacement recombination events between the native sequence and any linear DNA fragment flanked by short (>40nt) homology arms (Figure 1). In contrast, this sort of sequence modifications cannot be done with our native excisable antibiotic-based system where one recombineering cassette needs to be constructed for each new tag. In order to bypass this limitation and, at the same time, to further facilitate the generation of new recombineering cassettes, we have developed two new recombineering cassettes, a *Universal tag-generator* cassette (where the counter-selectable marker *RPSL* allows for the selection of DNA replacement events in the presence of streptomycin) and a *galK-FRT-Amp-FRT* cassette (where *galK* can be employed as a contra-selectable marker) (Figure 2, Table S5). These two cassettes can be used to facilitate the addition of new tags to our collection of bifunctional recombineering cassettes by simply replacing the *RPSL* or the *galK* sequences by the sequence of a new tag, or to generate nearly any types of gene editing events, from single nucleotide modifications to large deletions, by replacing the whole cassette by the sequence of interest by recombineering (Figure 2) (Stepanova et al., 2011; Brumos et al., 2018). As a proof of concept, we have used the *Universal tag-generator* cassette to create a new *RFP* recombineering cassette and the *galK-FTR-Amp-FRT* to generate the *GFP, mCherry* and *3xMYC* recombineering cassettes (Table S5).

**Figure 2.**
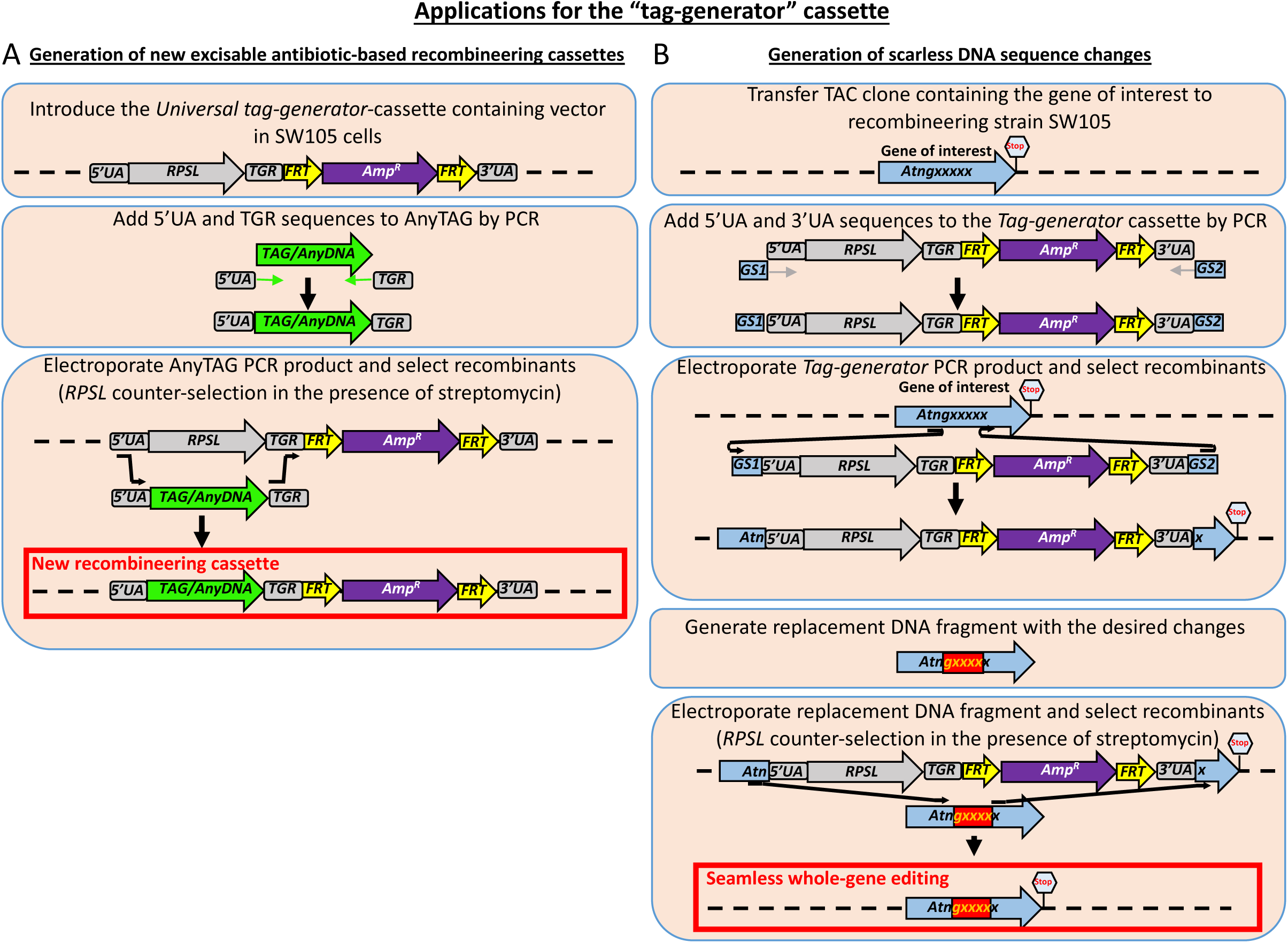
Schematic representation of two applications for the *tag-generator* cassette. A *tag-generator* cassette consisting of the negative selectable marker gene *RPSL* and a positive selectable marker *Amp^R^* conferring ampicillin resistance can be used for the easy generation of new bifunctional recombineering cassettes containing any desired tag **(A)**, or to make precise gene editing (such as introducing point mutations, deletions, or insertions) in the gene of interest **(B)**. To facilitate the use of this *tag-generator* cassette, in addition to the negative (*RPSL*) and positive (*Amp^R^*) selectable markers, the construct contains the 5’ and 3’ universal adaptors (UA) that allow for the amplification of any recombineering cassette in our collection (see below) and the *TGR* sequence that allows for the in-frame insertion of any tag, making it possible to use the resulting cassettes in tagging experiments at any position in the gene of interest (N-terminal, C-terminal, or internal). Finally, this cassette also includes *FRT* sites flanking the sequences conferring ampicillin resistance (*Amp^R^*) allowing for the precise and efficient elimination of the selectable marker gene post-insertion. The *tag-generator* cassette can be used to construct new recombineering cassettes. **(A)** A ready-to-use SW105 *E. coli* strain containing a TAC clone that harbors the *tag-generator* cassette has been constructed (top panel). Using the primers TGF (5’-TAAAAAGGGTTCTCGTTGCTAAGGAGGTGGAGGTGGAGCT-3’ in-frame with 20 nucleotides of the 5’ of the new tag) and TGR (5’-gaaagtataggaacttcccacctgcagctccacctgcagc-3’ in frame with 20 nucleotides that anneal to the 3’ end of the tag of interest), the tag of interest (*TAG*/*AnyDNA*) can be amplified generating the *5’UA-TAG/AnyDNA-TGR* amplicon (middle panel). By electroporating this amplicon in the *SW105* recombineering strain carrying the *tag-generator* cassette and selecting for the absence of *RPSL* (streptomycin-resistant colonies), a new bifunctional recombineering cassette for the tag of interest will be obtained (bottom panel). **(B)** The tag generator cassette can also be used in a two-step recombination procedure similar to the classical *galK* approach to make any type of sequence modification, such as seamless insertion of a tag, introduction of point mutations, etc. In this case, the process starts with the identification of the genomic clone containing the gene of interest (top panel). Using GS1 and GS2 primers (see figure 1) to PCR-amplify the *tag-generator* cassette, an amplicon containing the sequences flanking the point where the gene editing will take place is obtained (second panel). By electroporating this amplicon in *SW105* recombineering cells carrying the BAC or TAC clone with the desired gene and selecting for ampicillin-resistant colonies, the gene of interest is tagged with the *tag-generator* cassette (third panel). Next, a replacement DNA construct containing the edited sequence (point mutations, deletions, insertions, etc. depicted as a red box in the fourth panel) flanked by long regions of homology to the gene of interest (100 to 200 base pairs on each side of the region to be edited are recommended) is produced, typically, by commercial DNA synthesis. When designing these constructs, it is important to consider that recombination can take place at any point within the regions of homology between the replacement sequence and the gene of interest tagged with the *tag-generator* cassette (bottom panel). By electroporating the replacement DNA and selecting for colonies resistant to streptomycin, the desired final product is obtained (bottom panel).

### Recombineering-based trimming and transfer of large genomic constructs from BACs to binary vectors

As indicated above, the ability to precisely edit the sequence of a GOI in the context of a large BAC has the great advantage of capturing distant (even tens of thousands of base pair away) regulatory sequences and, thus, preserving the native expression patterns in the transgene reporter fusions. The use of BACs containing the GOI as the source of the genomic sequences to be edited has, however, several critical drawbacks. First, the researcher does not have the flexibility to choose the exact DNA regions flanking the GOI that would be included in the final construct, as this would be determined by the sequences already present in the selected BAC clone. Second, the choice of sequence-indexed BACs containing the GOI is limited by what clones are available in the BAC collection. Additionally, in most plant species (with probably the sole exception of *Arabidopsis*), the BAC clone collections that have been mapped back to the genome cannot be directly used for *Agrobacterium*-mediated transformation, as the vectors used in various genome sequencing efforts lack the features for propagation in *Agrobacterium* and for the subsequent transfer of DNA from the bacteria to the plant genome. To circumvent these limitations, we have developed a set of antibiotic-selection-based recombineering “trimming cassettes” (Figure 3, Table S5) that allow for the efficient elimination of undesired sequences flanking the GOI (Figure 3A). This simple trimming procedure allows the researcher to precisely define the DNA regions flanking the GOI to be included in the final construct (assuming a BAC clone containing the desired regions has been identified), thus eliminating extra genes that may cause phenotypic alterations when present in a copy-number excess. An added advantage of this strategy is that by reducing the size of the final construct, the transferring efficiency of the desired edited sequences to the plant nuclear genome is also increased (Zhou et al., 2011; Brumos et al., 2018).

**Figure 3.**
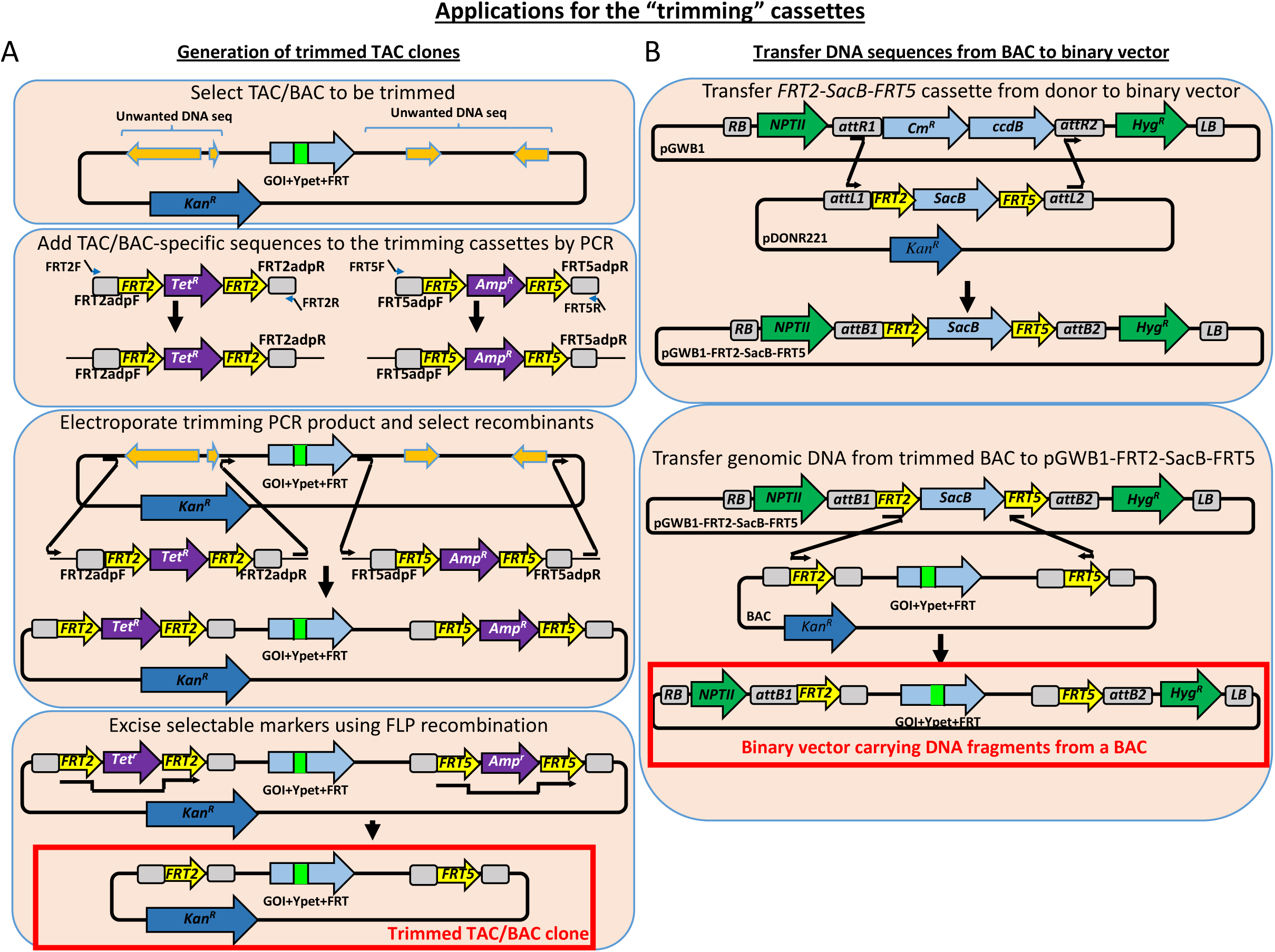
Schematic representation of two applications for the trimming cassettes. Two trimming cassettes, one conferring tetracycline resistance and another conferring ampicillin resistance, have been generated to facilitate the elimination of undesired sequences in TAC or BAC clones, as well as for the efficient transfer of large fragments of DNA from BAC clones to binary vectors. To make these actions possible, each antibiotic selectable marker in the trimming cassettes is flanked by a different pair of orthogonal *FRT* sequences, *FRT2* or *FRT5*, that not only allow for the elimination of the antibiotic-resistance sequences after the trimming process **(A)**, but also for the efficient *in vivo* transfer of large fragments of DNA from a BAC or TAC clone to a modified binary vector **(B)**. **(A)** The first step in the process of trimming a genomic sequence is to identify a BAC or TAC clone carrying the gene of interest (top panel). Using DNA for the ready-to-use *FRT2-Tet-FRT2* and *FRT5-Amp-FRT5* trimming cassettes as PCR templates and two pairs of primers, FRT2F/FRT2R, and FRT5F/FRT5R (see sequences characteristics below), two amplicons containing the sequences of the trimming cassettes flanked by 40 nucleotides homologous to the sequences flanking the region to be deleted in the target genomic DNA are produced by PCR (second panel). Electroporating these amplicons into electrocompetent SW105 cells carrying the TAC clone harboring the gene of interest and selecting for colonies resistant to both ampicillin and tetracycline results in the replacement of the undesired genomic DNA sequences by the trimming cassette sequences (third panel). Inducing the expression of the *FLP* recombinase present in the genome of the SW105 cells results in the elimination of the ampicillin and tetracycline selectable sequences, leaving behind a single *FRT2* and *FRT5* site at each flank, respectively (bottom panel). **(B)** The trimming product obtained in (A) contains the desired genomic DNA fragment flanked by two orthogonal *FRT* sites opening the possibility of using cassette-exchange strategies to move this potentially large DNA from the original BAC/TAC to a binary vector. To generate binary vectors suitable for this cassette-exchange reaction, we first generated a derivative of the Gateway pDONR221 vector containing the negative selectable marker *SacB* flanked by the head-to-toe *FRT2* and *FRT5* sites (top panel). Using this new vector, the *FRT2-SacB-FRT5* cassette can be easily transferred to any attR1-attR2-containing destination vector such as pGWB1 (top panel). To transfer the genomic DNA fragment flanked by the *FRT2* and *FRT5* sites to the pGWB1-FRT2-SacB-FRT5 vector, SW105 cells carrying the trimmed BAC or TAC clone (from bottom panel in (A)) can be electroporated with the pGWB1-FRT2-SacB-FRT5 vector. In the presence of sucrose (negative selection for the *SacB* gene) and hygromycin (positive selection for the pGWB1 backbone), the product of a successful cassette-exchange reaction can be efficiently selected. Dark green arrows indicate resistant genes that work both in plants and bacteria. The primers used to amplify the trimming cassettes have the following structure. FRT2 F: 5’-40nt corresponding to the sequence upstream of the nucleotide in front of which one wants to insert the *FRT2* site followed by the sequence -ttcaaatatgtatccgctca -3’. FRT2 R: 5’- 40nt corresponding to the reverse complement sequence downstream of the nucleotide after which one wants to insert the *FRT2* site followed by the sequence -ttaccaatgcttaatcagtg -3’. FRT5 F: 5’-40nt corresponding to the sequence upstream of the nucleotide in front of which one wants to insert the *FRT5* site-aacgaatgctagtctagctg-3’. FRT5 R: 5’-40nt corresponding to the reverse complement sequence downstream of the nucleotide after which one wants to insert the *FRT5* site-ttagttgactgtcagctgtc -3’.

In addition to the antibiotic resistance markers present in these trimming cassettes, we have also included two sets of orthogonal *FRT* sites (*FRT2* in the tetracycline and *FRT5* in the ampicillin cassettes, respectively (Schlake and Bode, 1994)), thus allowing for the removal of the antibiotic resistance genes once the trimming has been completed. Importantly, after the *FLP*-mediated excision of the antibiotic resistance genes, the two remaining *FRT2* and *FRT5* sites left in the construct display a head-to-tail orientation. As illustrated in Figure 3B and S1, this *FRT* configuration allows for the transfer of the selected DNA flanked by the *FRT*s to an engineered binary vector (see below) through an *in vivo* cassette-exchange reaction ((Turan et al., 2013) and Figure 3B and S1). To carry out the transfer of BAC DNA to any Gateway-compatible binary vector, we have constructed a pDONR221-based entry clone with the negative selectable marker *SacB* flanked by the *FRT2* and *FRT5* sites in the same head-to-tail configuration as in the trimmed BAC (Figure 3B, Table S5). This *FRT2-SacB-FRT5* cassette can now be transferred to any *attR1-attR2*-containing destination vector using the standard LR Gateway recombination system, making it capable of accepting an *FRT2/FRT5*-flanked insert from any BAC clone.

One possible advantage of this *in vivo FLP*-based cassette exchange system relative to the *in vitro* systems such as Gateway is the higher upper size limit of DNA fragments that can be routinely mobilized between vectors. Following this strategy, we have generated a pGWB1-FRT2-SacB-FRT5 as a standard destination vector for our *FLP*-mediated cassette exchange reactions (Figure 3B, Table S5). To test the efficiency of the *FLP*-based system to exchange large DNA fragments, we tested the ability to transfer DNA fragments of ∼16, ∼37 and ∼78 kb from a BAC containing the *YUC9-GUS* translation fusion gene to the pGWB1-FRT2-SacB-FRT5 binary vector. Although we were able to transfer all three DNA fragments, we found that the efficiency of the transfer dropped considerably as the DNA fragment size increased (Table 1). We reasoned that this could be due to a compromised stability of very large constructs in a multicopy plasmid such as pGWB1 not designed to hold such large DNA inserts. To overcome such limitation, we engineered pYLTAC17, a low-copy vector designed for the generation of large-insert genomic TAC libraries (Liu et al., 2002), to carry the exchange cassette *FRT2-SacB-FRT5*, allowing for the transfer, stable propagation, and plant transformation of large fragments of DNA originally carried in a BAC clone (Figure S1, Table S5). Furthermore, to expand the spectrum of BAC libraries that can be used as a DNA donor in this system, we introduced *aadA*, an *aminoglycoside 3’-adenylyltransferase* gene that confers spectinomycin and streptomycin resistance in both in *E. coli* and *Agrobacterium*, in addition to the kanamycin-resistance gene already present in the pYLTAC17-FRT2-SacB-FRT5-Spect vector (see Methods) (Figure S1, Table S5). We have also generated a second version of this vector pYLTAC17-FRT2-SacB-FRT5-Spect-Kan where the *Bar* gene for phosphinothricin (Basta) resistance has been replaced by the *NPTII* gene for kanamycin selection in planta (Figure S1, Table S5). Using the pYLTAC17-FRT2-SacB-FRT5-Spect plasmid side-by-side with the pGWB1-FRT2-SacB-FRT5 vector, we observed in two independent experiments (see Table 1) that the efficiency of DNA transfer from the BAC to pYLTAC17-FRT2-SacB-FRT5-Spect was higher than that to pGWB1-FRT2-SacB-FRT5-Spect as the acceptor vector, especially for DNA fragments as large as 78kb.

**Table 1.** Efficiency of DNA transfer from the BAC IGF F20D22 to pGWB1-FRT2-SacB-FRT5 vectors and pYLTAC17-FRT2-SacB-FRT5-Spec-Kan

|  |  | pGWB1-FRT2-SacB-FRT5 |  |  | pYLTAC17-FRT2-SacB-FRT5-Spec-Kan |  |  |
| --- | --- | --- | --- | --- | --- | --- | --- |
|  |  | Colony number for PCR | Positive colonies | % | Colony number for PCR | Positive colonies | % |
| 1<sup>st</sup> experiment | JMA2364 (~16 kb) | 10 | 8 | 80 | 10 | 10 | 100 |
|  | JMA2365 (~37 kb) | 7 | 5 | 71.4 | 10 | 10 | 100 |
|  | JMA2366 (~78 kb) | 3 | 1 | 33.3 | 10 | 10 | 100 |
| 2<sup>nd</sup> experiment | JMA2364 (~16 kb) | 10 | 10 | 100 | 10 | 10 | 100 |
|  | JMA2365 (~37 kb) | 10 | 9 | 90 | 10 | 10 | 100 |
|  | JMA2366 (~78 kb) | 6 | 3 | 50 | 10 | 9 | 90 |

### High-throughput recombineering using highly efficient FLP-based marker excision cassettes

In the post-genome era, with thousands of gene sequences available, scalability represents a key element of any experimental procedure that aims to facilitate gene functional analysis. With the goal of developing a simple pipeline to process 96 recombineering samples in parallel (Figure 4 and Methods), different bottlenecks were identified. The first challenge was the development of an efficient method to transfer 96 TAC clones from the original *E. coli* strain DH10B to the recombineering strain of *E. coli*, SW105. This problem was addressed by growing the 96 DH10B strains in a 96-deep-well plate overnight and carrying out standard home-made alkaline lysis miniprep (Alonso and Stepanova, 2014) in a 96-well format. Critical to the robustness of this procedure was the gentle manipulation of the TAC DNA (e.g., no vortexing or freezing) to avoid breaks or nicks that would result in the loss of its supercoiled conformation and a drastic decrease in its transformation efficiency. Towards the same objective of maintaining the supercoiled conformation of the TAC DNA, electroporation was carried out immediately after the DNA purification procedure was finished. Electrocompetent SW105 cells were freshly prepared using standard procedures and electroporation carried out using a 96-well-format electroporator (Alonso and Stepanova, 2014). Cells corresponding to individual clones were then plated and individual colonies tested for the presence of the correct TAC clone by PCR using gene-specific primers. A similar strategy was used to transfer the TAC clones from *E. coli* SW105 to the *recA-Agrobacterium* UIA143 pMP90 once the desired modifications had been introduced into the genes of interest and the constructs confirmed by PCR and sequencing. The next critical step that needed to be scaled up was the insertion of the tag in the desired locations in each of the 96 selected genes. PCRs with a 60mer primer pair containing 40 nucleotides flanking the insertion site of the GOI and 20 nucleotides corresponding to the universal adaptors flanking the recombineering cassette were used to obtain the 96 gene-indexed recombineering amplicons. Key for the implementation of this high-throughput procedure was the experimental design of a strategy that would allow for an efficient introduction of 96 different recombineering cassettes into 96 different SW105 strains carrying the individual BACs of interest without the need to having to individually prepare electrocompetent cells for each of the 96 SW105 strains. This was achieved by preparing electrocompetent cells from pools of 12 strains corresponding to a full row in the 96-well plate in such a way that 96 TAC clones were represented in 8 non-redundant pools of competent cells per plate. Each of these pools of competent cells was divided in 12 identical aliquots, with each aliquot electroporated with one of the 12 amplicons containing the 40 nt flanking sequences specific for one of the 12 targets contained in this pool (Figure 4). Due to the high sequence specificity of the recombination events, only those cells in a pool carrying the gene corresponding to the particular gene-indexed recombineering amplicon can undergo recombination and, therefore, acquire the selectable marker encoded in the cassette. For each of the 96 parallel recombination experiments, the fidelity of the recombination events was assayed by PCR using gene-specific primers flanking the selected insertion site with an efficiency of ∼100%, as we have previously reported (Zhou et al., 2011). To excise the selectable marker, the 96 strains confirmed to carry the desired gene fusions were grown overnight in a 96-deep-well plate. Cells from each well were then diluted 50-fold in 1 mL of fresh LB media in a new 96-deep-well plate and the *FLP* gene was induced by adding 10 µL of 10% (w/v) L-arabinose per well. After 2h at 32°C, a sterile toothpick was used to streak a few cells in a solid media plate. For the first 96 constructs, we used the recombineering cassette containing the *galK* marker flanked by the *FRT* sites with the idea that the positive/negative selection of *galK* could be used to select for galK-clones after the *FLP*-mediated excision. Due to the extremely high efficiency of excision, we found the *galK* contra-selection unnecessary, as the desired excision events for most clones (see below) could be identified without the need for contra-selection. In fact, the analysis of three independent clones for each construct was sufficient in most cases to find at least one excision event lacking any undesired mutation (see below).

**Figure 4.**
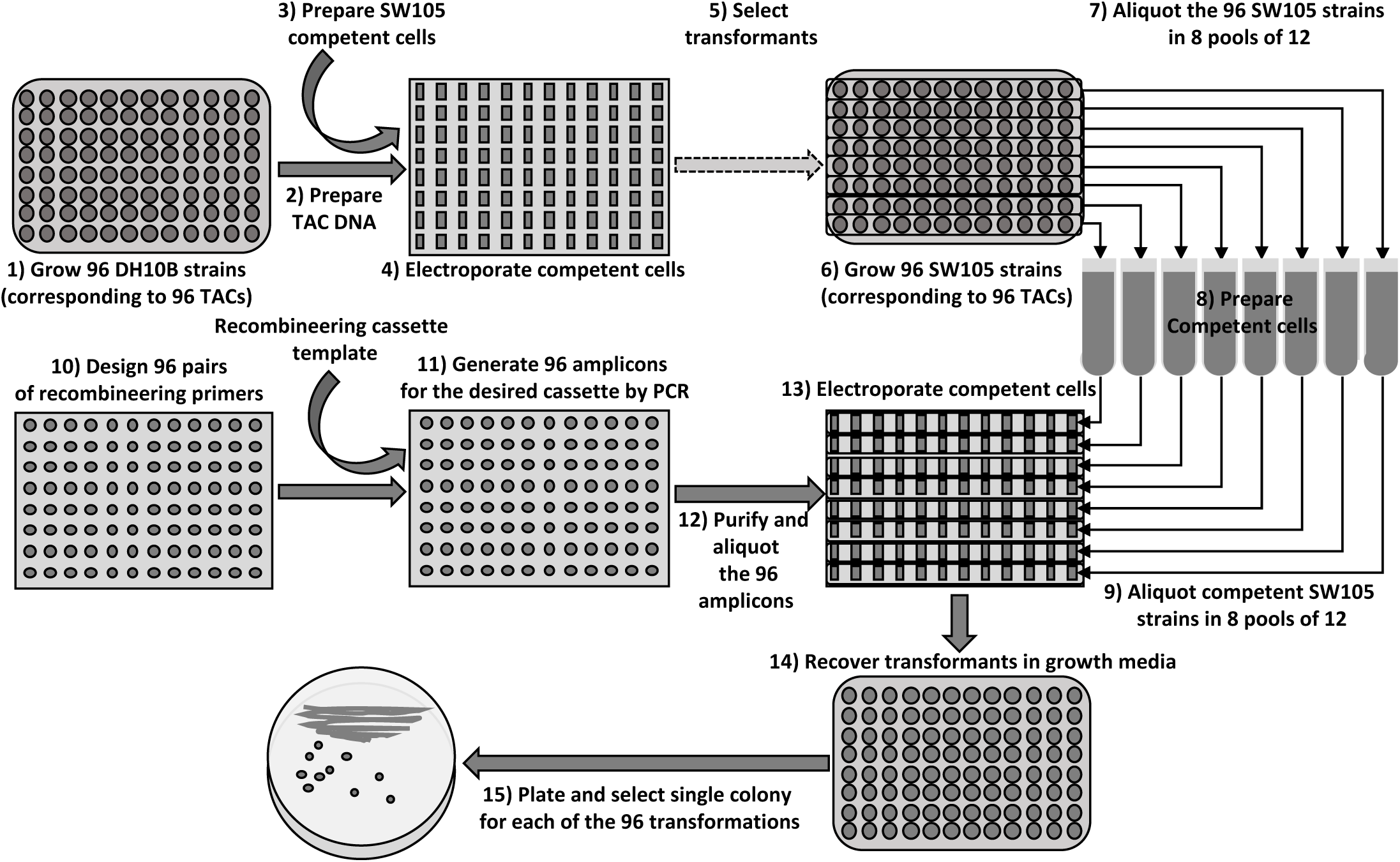
Schematic representation of the high-throughput recombineering pipeline. The process starts by growing 96 DH10B strains carrying the desired TAC clones (best TAC clones from the two available *Arabidopsis* libraries for any given gene can be found in our genome browser at https://brcwebportal.cos.ncsu.edu/plant-riboprints/ArabidopsisJBrowser) in a 96-deep-well plate (1). The cells are pelleted by centrifugation and a 96-well-format alkaline-lysis DNA miniprep protocol is used to obtain DNA for the corresponding 96 TACs (2). Electrocompetent SW105 cells are prepared and aliquoted into a 96-well electroporator cuvette (3). DNA for each of the selected 96 TAC clones is added to the electroporation cuvette wells and electroporated into the SW105 competent cells (4). After the electroporation, cells are resuspended in LB and transferred to a 96-deep-well plate where they are allowed to recover before they are plated in selectable media. Individual clones grown in the selectable media are tested by PCR and arranged back into a 96-well format (dashed arrow indicates that several steps are not shown) (5). The SW105 strains carrying 96 TAC clones selected in step 5 are grown overnight in a 96-deep-well plate (6). Cells from the overnight culture are used to inoculate 8 cultures corresponding to pools of 12 clones each (7). Electrocompetent cells from each of the 8 pools of 12 clones are prepared (8). Aliquots of cells from each pool are placed into the wells of the corresponding rows of the 96-well electroporation cuvette. For example, from pool one, 12 identical aliquots would be placed in each of the wells of the first row of the 96-well electroporator cuvette and so on (9). In parallel, a pair of 60mers per gene are designed (primer sequences for generating N- and C-terminal amplicons for any gene and any of our ready-to-use recombineering cassettes can be obtained from our genome browser at https://brcwebportal.cos.ncsu.edu/plant-riboprints/ArabidopsisJBrowser) (10) and used to generate the corresponding 96 amplicons using the DNA from one of our ready-to-use cassettes as a template (11). The amplicons are purified by simple chloroform extraction and ethanol precipitation in a 96-well plate (12). The corresponding 96 amplicons are added to the electrocompetent cells and electroporated in the 96-well electroporation cuvette (13). As before, the cells are resuspended in LB and transferred to a 96-deep-well plate to allow them to recover (14). The cells from each transformation are then streaked in LB plates with the proper antibiotic (15). Individual colonies (one or two per construct) are examined by colony PCR using a combination of gene- and tag-specific primers and the integrity and fidelity of the recombination is checked by PCR fragment sequencing.

As indicated above, the first 96 genes (Table S1) corresponding to hormone-related genes were tagged using a modified version of a previously developed (Tursun et al., 2009) *Venus-FRT-galK-FRT* cassette (Table S5) where we added the universal adapters mentioned above (Tian et al., 2004). From these 96 selected genes, we failed to generate the desired constructs in two cases where, after sequencing three independent clones, we were not able to identify a construct with the desired modifications. For 17 additional genes we had to sequence two clones to find the desired mutation-free construct, and in three cases a third clone had to be sequenced. As we have previously described (Zhou et al., 2011), most of the observed mutations were found in the sequences corresponding to the long oligos used to amplify the recombineering cassettes. After sequence verification, 80 out of 94 clones were successfully transferred to a *recA-Agrobacterium* strain UIA143 pMP90 (Hamilton, 1997) using our 96-well-plate pipeline described above (Figure 4 and Methods). In the 14 cases we did not succeed to transfer the TAC clone to *Agrobacterium*, we did observe *Agrobacterium* colonies growing in kanamycin-selectable media, but they tested negative for the presence of the tagged gene by PCR (see below).

Importantly, as mentioned above, we observed that the efficiency of *FLP*-based excision of the *galK* cassette was ∼100% efficient, even in the absence of counter-selection conditions, indicating that the positive/negative selectable marker *galK* could be replaced by a much more convenient antibiotic-based positive-only selection marker, allowing for the use of standard growth media (instead of the minimal media required in the *galK* system). In addition to lowering the complexity and cost of the recombineering experiments, the use of antibiotic-based cassettes also reduces significantly the time required for *E.coli* to grow in the selectable media, going from 5 days for the *galK* selection in M63 minimal media to 2 days (as the recombineering strains need to be grown at 32°C to avoid the induction of the lambda red proteins) for the antibiotic-based selection in standard LB media (Figure 1). To test the utility of these antibiotic-based recombineering cassettes, we generated the *Universal AraYpet-FRT-Amp-FRT* cassette and used it to tag another set of 96 genes (Table S5 and S2). In this second experiment, we included most of the genes in the shikimate and shikimate-derived metabolic pathways, focusing on those related to auxin biosynthesis. Similar to the *galK*-based system, we were able to obtain mutation-free constructs for most of the genes (89 out of 96) and transfer them to *Agrobacterium* in 79 out of the 89 cases. Although we are not sure what the problem was in the 12 cases where the *Agrobacterium* transformation failed, in a follow-up study we have found that by adding *aadA* (Sandvang, 1999) as a second antibiotic selectable marker we can eliminate false positives during the transfer of large TAC clones from *E. coli* to *Agrobacterium* (see below), thus improving the efficiency of selection of TAC clones in *Agrobacterium* to ∼100%.

All 159 *Venus* or *Ypet* constructs transferred to *Agrobacterium* were used to transform *Arabidopsis* using the highly efficient floral dip method but replacing the sucrose by glucose to prevent toxicity in some *Agrobacterium* strain where the *SacB* gene was still active (Zhou et al., 2011). To facilitate the plant transformation process of the large number of constructs generated, the 159 *Agrobacterium* strains were grown in solid media (two 150 mm Petri dishes per construct) and *Agrobacterium* cells were then harvested in transformation media just before performing the floral dip (see Methods for more details). Of these 159 constructs, we have generated and deposited in the stock center *Arabidopsis* transgenic lines for 33 genes (two independent lines for 31 of these genes and one single line for the other two) (see Tables S1-S3 for accession number information). This subset of lines was selected based on an initial screen of young T1 seedlings with positive fluorescence signal and subsequent PCR confirmation of the desired genotype. We decided to prioritize this relatively small subset of genes due to the resources that would be needed for (and the logistic challenges that would be involved in) the propagation, making homozygous, and subsequent characterization of several lines per construct for which no evidence of detectable fluorescence and, therefore, future utility was readily available. The lack of detectable expression of the reporter gene could be due to several factors. On the one hand, we have observed that rates of deletions of the TAC constructs during the plant transformation process could be quite significant for large constructs, while negligible for constructs smaller than 25kb (Zhou et al., 2011). Additional and probably a more significant factor is the low expression/accumulation levels of many of the tagged proteins. To offset the first problem, we could either identify by PCR transgenic lines containing the whole transgene including both ends of the T-DNA, as we have done previously (Zhou et al., 2011), or we could trim distal genomic sequences unlikely to contain regulatory elements affecting the expression of the GOI but present in the original TAC clones. Much more difficult to circumvent is the problem of lack of detectable fluorescence signal due to low levels of expression. To try to alleviate these two problems derived from using large TAC clones and weak florescence signal from low expressed genes, we selected 87 of previously tagged genes related to auxin biosynthesis, transport and response and generated new recombineering constructs tagged with three copies of the bright fluorescent protein gene *Ypet* (Table S5, and S3). Towards this end, we made a new codon-optimized, *FLP*-based, ampicillin-resistant, excisable recombineering cassette (Table S5). At the same time, all these new constructs were also trimmed to reduce the insert to just the tagged gene and 15kb of flanking sequences (10kb upstream of ATG and 5 kb downstream of the stop codon) using the trimming tools described above.

### Characterization of expression patterns for *TAA1/TAR* and *YUC* genes

To further demonstrate the utility of this high-throughput recombineering system, we employed a new codon-optimized *GUS* recombineering cassette to tag the 14 auxin biosynthetic genes of the IPyA pathway (Table S3): *TAA1, TAR1, TAR2*, and *YUC1* to *YUC11*. *TAA1* and *TAR*s encode tryptophan aminotransferases that catalyze the synthesis of IPyA from tryptophan (Stepanova et al., 2008; Tao et al., 2008), whereas *YUC1* to *YUC11* are flavin monooxygenases that convert IPyA to IAA (Sugawara et al., 2009; Stepanova et al., 2011). We generated transgenic plants for all 14 genes and examined expression patterns of translational fusions in seedlings and reproductive tissues (Figures 5, 6, 7).

In roots of three-day-old dark-grown seedlings germinated in control AT media, we could detect the expression of translational fusions with *GUS* for *TAA1* and four out of eleven *YUC*s (*YUC3, YUC7, YUC8*, and *YUC9*) in the primary root meristem, as well as *TAA1* and *YUC6* in the pre-vasculature (Figure 5). Treatments with the auxin transport inhibitor naphthylphthalamic acid (10 µM NPA), ethylene precursor 1-aminocyclopropane-1-carboxylic acid (10 µM ACC), NPA and ACC combined, or synthetic auxin naphthaleneacetic acid (50 nM NAA) enabled the detection of root expression of all three *TAA1/TAR* genes and nine out of eleven *YUC*s, except for *YUC1* and *YUC10* that were not expressed in distal regions of the primary root of three-day-old etiolated plants in any of the conditions tested. ACC treatment upregulated *TAA1, TAR1, TAR2, YUC2, YUC3, YUC4, YUC5, YUC6, YUC8, YUC9*, and *YUC11*, consistent with the induction of the auxin responsive reporter *DR5:GUS* (Figure 5) and the known stimulatory effect of ethylene on auxin biosynthesis in roots (Stepanova et al., 2005; Ruzicka et al., 2007; Stepanova et al., 2007; Swarup et al., 2007). Germinating seedlings in the presence of NPA induced the levels of *TAA1, TAR2, YUC3, YUC5, YUC7, YUC8, YUC11*, and, accordingly, *DR5*, but the domains of NPA-triggered GUS activity were different for different genes. For example, for *TAA1* and *YUC5*, GUS staining in NPA was visible in the root elongation zones, suggesting that local auxin production is activated in this part of the root in response to the inhibition of polar auxin transport. Furthermore, the expression of *TAA1* in the developing vasculature and of *TAR2* in the stele was also enhanced by NPA. The domains of *YUC3* and *YUC8* in NPA became dramatically expanded in the primary root meristems, presumably leading to increased local production and accumulation of auxin in these tissues, as witnessed by the extensive widening of the *DR5:GUS* domains. The shift of the *DR5* maximum correlates with the previously reported broadening of the stem cell niche under NPA treatments (Sabatini et al., 1999). The re-patterning of the meristematic tissues is triggered by the increased levels of IAA trapped in the auxin-producer cells, with similar outcomes described for root meristems in plants exposed to an exogenous synthetic auxin, 2,4-D (Sabatini et al., 1999).

**Figure 5.**
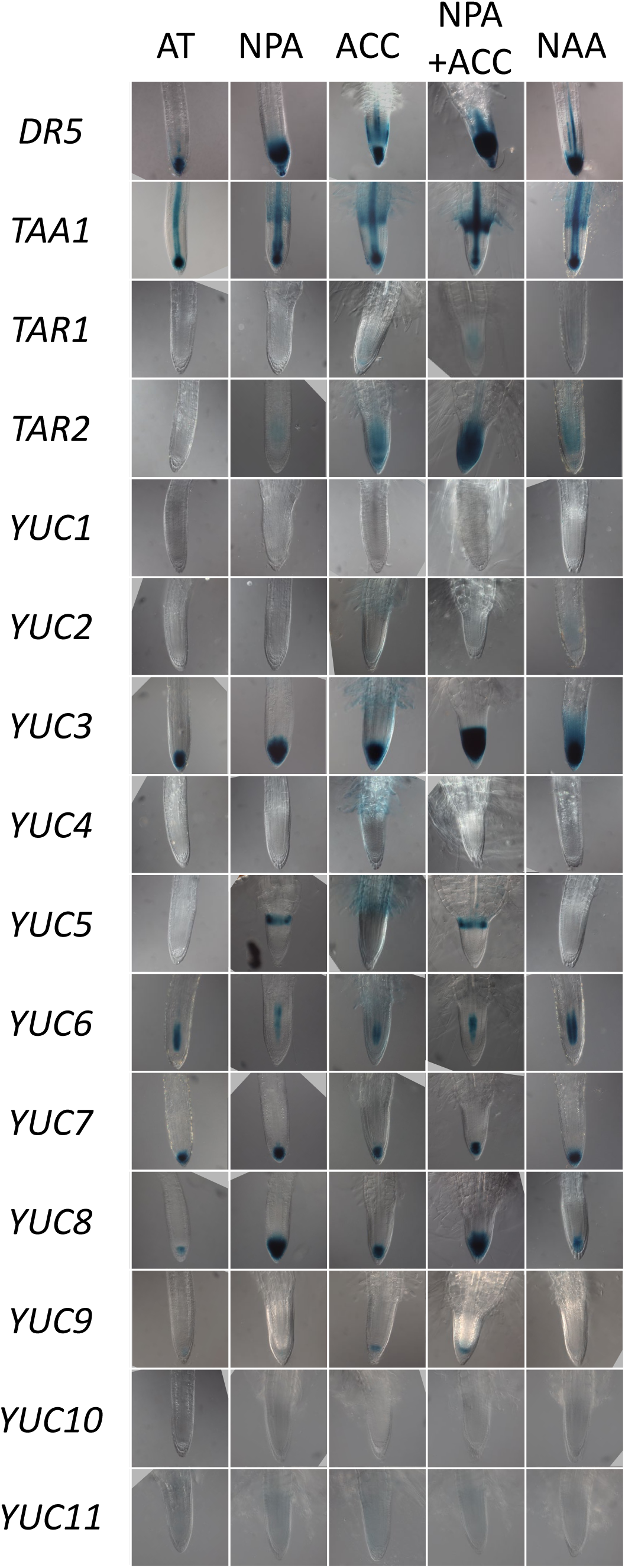
GUS staining patterns of translational recombineering fusions of auxin biosynthesis genes and *DR5:GUS* in roots. Seedlings were germinated for three days in the dark in control AT media or in AT media supplemented with 10uM NPA, 10uM ACC, 10uM NPA + 10uM ACC, or 50nM NAA. Samples were optically cleared with ClearSee.

Combined NPA plus ACC treatment had additive or synergistic effects on the expression of *TAA1, TAR1, TAR2, YUC3*, and *YUC9* or, in the case of *YUC5, YUC6, YUC8*, phenocopied the single NPA treatments (Figure 5). Interestingly, in some cases, combined NPA plus ACC treatment resulted in the loss of some of the subdomains of expression visible with ACC alone (e.g., GUS staining in root hairs for *YUC2, YUC3, YUC4, YUC5* and *YUC6*), or led to the shift in the domain of GUS activity, as seen for *TAR1*. Finally, the NAA treatment upregulated *TAA1* in the root elongation zone*, TAR2* and *YUC2* in the stele and root cap*, YUC3* in the entire root tip, *YUC6* in the vasculature, and *DR5:GUS* in the vasculature and the root meristem, suggesting that exogenous auxin can activate endogenous auxin biosynthesis. Of the 12 genes detectable in roots, only *YUC7* was not prominently responsive to any of the treatments tested (Figure 5).

In shoots of three-day-old etiolated seedlings, *TAA1, TAR2* and five *YUC* genes, *YUC1, YUC3, YUC4, YUC5*, and *YUC6*, were expressed in control media, whereas *TAR1, YUC2, YUC7, YUC8, YUC10* and *YUC11* became detectable in seedlings exposed to ACC, NPA plus ACC, and/or NAA (Figure 6). The spatial domains of *GUS* reporter activity varied for different auxin biosynthesis genes. For example, in control conditions, *TAA1, YUC1, YUC4* and, to a lower extent, *TAR2* had defined expression in the shoot apical meristem, *TAA1* and *YUC6* were active in the hypocotyl vasculature, whereas *TAA1*, *TAR2* and *YUC6* had some activity in the cotyledon vasculature. *YUC4* and *YUC5* showed complementary expression patterns along the cotyledon perimeter. *YUC4* expression concentrated in the distal end of the cotyledon and *YUC5* was active along the edge of the cotyledon without overlapping with the *YUC4* domain (Figure 6). These well-defined expression patterns of *GUS* fusions suggest that local auxin is produced in specific tissues by a combinatorial action of several tryptophan aminotransferases and flavin-containing monooxygenases that together contribute to establishing the morphogenic gradients of auxin.

**Figure 6.**
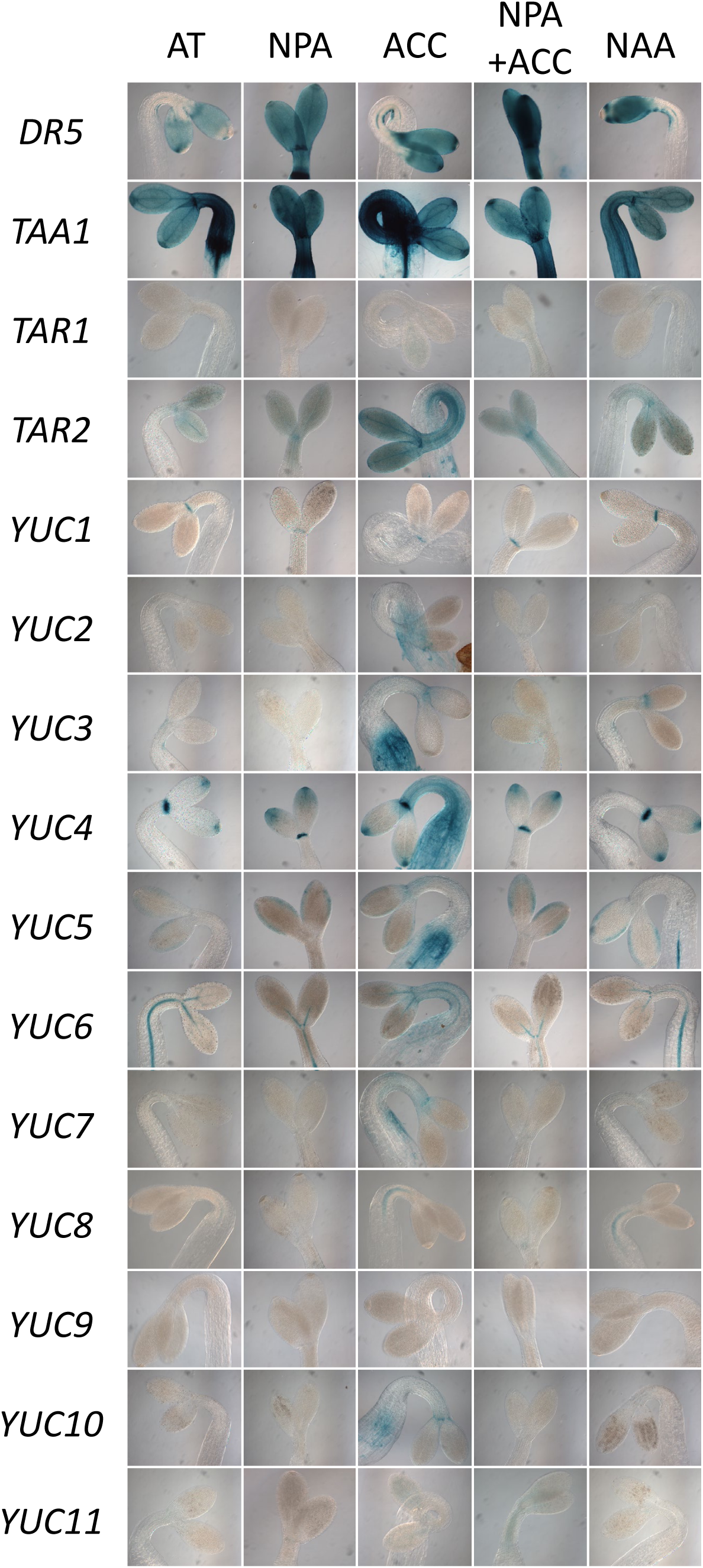
GUS staining patterns of translational recombineering fusions of auxin biosynthesis genes and *DR5:GUS* in shoots. Seedlings were germinated for three days in the dark in control AT media or in AT media supplemented with 10uM NPA, 10uM ACC, 10uM NPA + 10uM ACC, or 50nM NAA. Samples were optically cleared with ClearSee.

Of the pharmacological treatments tested in shoots of three-day-old etiolated seedlings, addition of ACC in the growth media had the greatest effect on auxin gene activity, inducing 10 of the 14 genes of the IPyA pathway, specifically *TAR2, YUC2, YUC3, YUC4, YUC5, YUC6, YUC7, YUC8, YUC10*, and *YUC11* (Figure 6). Remarkably, in the presence of NPA plus ACC, all of these nine genes showed patterns and levels of expression indistinguishable from that in NPA alone, indicating that NPA could block the effect of ACC in shoots and implying that ACC may exert its effect by inducing polar auxin transport, consistent with prior reports (Ruzicka et al., 2007; Swarup et al., 2007). In contrast, the poorly expressed *YUC11* displayed barely detectable activity in both ACC and in NPA plus ACC, but not in NPA alone. Of the 13 auxin biosynthesis genes detectable in shoots (all but *YUC9*), only *YUC1* was not notably responsive to any of the four pharmacological treatments (Figure 6).

We also tested the expression of the 14 auxin biosynthesis genes in inflorescences and flowers of soil-grown plants (Figure 7). *TAA1* showed predominant expression in young gynoecia, especially in the developing ovules, and somewhat milder expression in the transmitting tract and ovules of older gynoecia (red arrows in Figure 7). In young anthers, *TAA1* exhibited a broad domain of expression, but as the anthers matured, the domain of *TAA1* activity became more restricted, concentrating at the distal tips of these organs (red arrows in Figure 7). *TAR1* and *TAR2* were also expressed in the anthers of young flowers, and *TAR2* was additionally detectable in the gynoecium and in mature flowers’ petal and sepal abscission zones (red arrows in Figure 7). Complementing the expression of *TAR2* in the abscission zones of older flower organs were multiple *YUC* genes (all but *YUC9* and *YUC11*) (red arrows in Figure 7).

**Figure 7.**
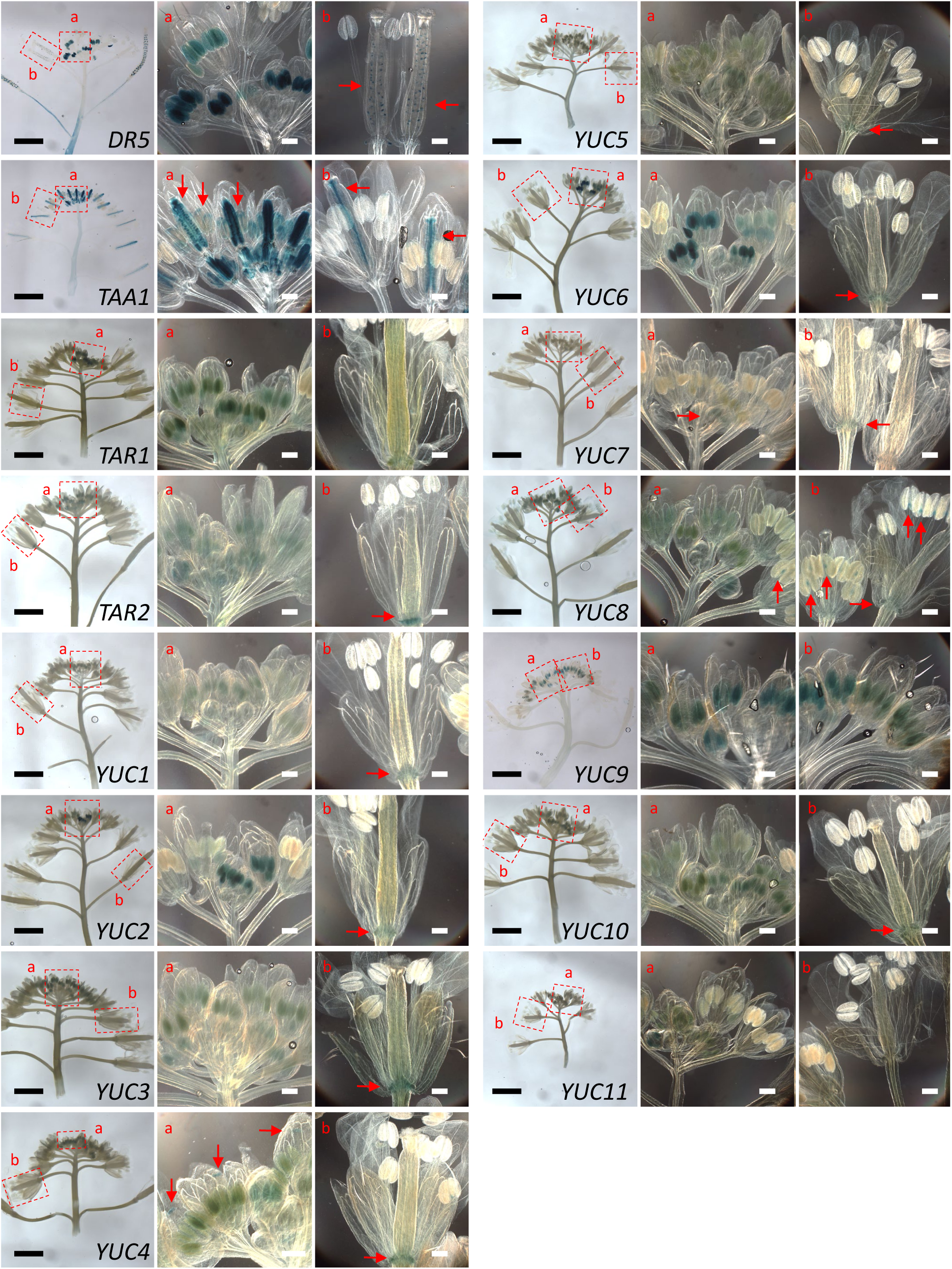
GUS staining patterns of translational recombineering fusions of auxin biosynthesis genes and *DR5:GUS* in inflorescences and flowers. Images of individual flowers represent the enlarged versions of the boxed areas of inflorescences. Red arrows mark the GUS activity domains of interest highlighted in the text. Black scale bars in the inflorescence images correspond to 2.5 mm. White scale bars in the flower pictures represent 250 mm. The samples of *DR5:GUS* and the *TAA1* recombineering fusion with *GUS* have been optically cleared with ClearSee to enable visualization of GUS activity in the ovules and developing seeds.

*DR5:GUS* and all members of the *YUC* family showed varying degree of activity in the anthers, with the immature male reproductive organs in *YUC2* and *YUC6* lines displaying the most prominent GUS activity, predominantly in the flowers of stage 8 to 13 (staging according to (Smyth et al., 1990; Alvarez-Buylla et al., 2010)). In older flowers, *YUC8* showed localized expression in the upper region of stamen filaments at the junctions with the anthers (red arrows in Figure 7). Remarkably, only *YUC4* was active in the gynoecia among all *YUC* family members (Villarino et al., 2016), specifically in the stigmatic tissue (red arrows in Figure 7). *DR5:GUS*, on the other hand, exhibited well-defined domains of expression in the ovules and developing seeds of older gynoecia (red arrows in Figure 7). None of the *YUC*s and *TAA1/TAR*s were prominently expressed in older anthers, petals or sepals (Figure 7). While we cannot exclude the possibility that some of the auxin biosynthesis genes are mildly active in those tissues, the expression levels of these enzyme genes fall below our detection limit.

## DISCUSSION

### Recombineering

High-efficiency homologous recombination mediated by the expression of specific phage proteins in bacteria, the process also known as recombineering, has been proven as an invaluable tool for high-throughput genome editing in bacteria (Isaacs et al., 2011). Although recombineering can equal and, in some respects, surpass the popular CRISPR-Cas systems as a precise genome editing tool in bacteria, to date, this system has not been proven to work efficiently in eukaryotic cells. Nevertheless, the power of recombineering has been widely used in eukaryotic model systems such as *C. elegans* (Sarov et al., 2006; Tursun et al., 2009) and *Drosophila* (Venken et al., 2008; Ejsmont et al., 2009; Sarov et al., 2016) to generate genome-wide collections of whole-gene translational fusions, thus opening doors to obtaining high-confidence gene expression landscapes in these organisms. As these whole-gene translational fusions are likely to capture most, if not all, of the regulatory sequences of a gene, it is not surprising that whenever systematic comparisons between classical and whole-gene recombineering-based translational fusions have been carried out, the superiority of the recombineering results has been clearly established (Sarov et al., 2012). Although no such systematic analysis has been carried out in plants, our anecdotal experience in *Arabidopsis* also suggests that recombineering-based whole-gene translational reporters are better at reflecting the native gene expression patterns. Thus, for example, the expression profiles of classical translational fusions for the auxin biosynthetic gene *TAA1* that passed the gold-standard quality control of complementing the mutant phenotype (Yamada et al., 2009) have been shown to be quite different from the expression domains observed with recombineering-based constructs (Stepanova et al., 2008). Importantly, we have recently shown that the recombineering-based, whole-gene, but not the classical translational fusion constructs, were able to complement a larger array of phenotypes examined under different conditions and in different tissues and mutant backgrounds (Brumos et al., 2018). Although somewhat anecdotal, the case of *TAA1* is not the only one reported, as the expression patterns deduced from an *AUX1* recombineering construct explain better than the classical promoter-fusion constructs the role of this gene in auxin redistribution in the root (Band et al., 2014). Furthermore, herein we show that the recombineering-construct-derived expression patterns of *TAA1* and several *YUC* genes (*YUC1*, *YUC2, YUC3, YUC4, YUC5*, and *YUC7*) are different from that previously reported using classical promoter fusions (Cheng et al., 2006; Yamada et al., 2009; Lee et al., 2012; Chen et al., 2014; Challa et al., 2016; Kasahara, 2016; Brumos et al., 2018; Xu et al., 2018) (see below). Again, although no systematic or comprehensive comparison has yet been performed in plants, the few examples described here in a plant with a relatively small and compact genome such as *Arabidopsis*, as well as the systematic analysis in *C. elegans*, strongly argue for the use of caution when inferring native expression patterns from translation-fusion experiments that use only a few kilobases of genomic DNA flanking the GOI. It is logical to think that the need for the use of large genomic regions flanking the GOI should be even greater in plants with larger (i.e., less compact) genomes. Ideally, direct tagging of the GOI in its chromosomal context should produce the most reliable expression patterns, but the current technology is not yet efficient enough to be widely adopted. It is likely that, in the same way that the constant advances in the CRISPR-Cas system technologies have made the introduction of mutations in a particular gene almost a routine in many plant research laboratories, the precise editing and insertion/replacement of sequences may also become habitual in the future. At this point, however, recombineering is the best alternative as it offers a relatively simple way to generating translational fusions and other types of gene editing events in the pseudo-chromosomal context of large bacterial artificial chromosomes. Nonetheless, to take full advantage of the power of recombineering, experimental-system-specific resources and tools need to be developed (Venken et al., 2006; Poser et al., 2008; Ejsmont et al., 2009; Tursun et al., 2009; Venken et al., 2009; Sarov et al., 2016).

In the past, we and others have shown that recombineering could be used to make precise gene modifications in the context of large DNA constructs in plants (Stepanova et al., 2008; Bitrian et al., 2011; Stepanova et al., 2011; Zhou et al., 2011; Peret et al., 2012a; Peret et al., 2012b; Pietra et al., 2013; Band et al., 2014; Fabregas et al., 2015; Han et al., 2015; Worden et al., 2015; Bhosale et al., 2018; Brumos et al., 2018; Yanagisawa et al., 2018; Gomez et al., 2019). In spite of the obvious advantages of using large fragments of DNA to ensure that most, if not all, regulatory sequences have been captured, and the relative ease by which different types of modifications can be introduced in large DNA clones such as BACs or TACs, recombineering has, at present, not been widely embraced by the plant community. Although there are probably several reasons for this, the extra labor and time required to generate recombineering constructs, the limited access to sequenced TAC libraries, the difficulty of working with large DNA constructs, etc. are among the likely factors.

To eliminate some of these potential obstacles for adopting recombineering and, thus, make this technology more accessible, we have developed and made freely available a new set of tools and resources. A collection of recombineering cassettes that contain both a commonly used tag (such as *GFP, GUS* etc.) and an antibiotic resistance marker have been generated (Table S5). In these cassettes, the sequences of the antibiotic resistance gene can be precisely removed with ∼100% efficiency using the *FRT* sites flanking the sequence by inducing a *FLP* recombinase integrated in the recombineering SW105 strain of *E. coli*. Using this new set of recombineering cassettes not only makes the procedure much faster and cheaper, but also extremely efficient and simple, all while avoiding the use of complicated and expensive bacterial minimal growth media. Limited access to transformation-ready bacterial artificial chromosomes containing the GOI could have also limited the adoption of this technology. To eliminate this potential problem, we have deposited in the ABRC a copy of the JAtY library developed at the John Innes Centre by Dr. Ian Bancroft‘s group. This, together with the recent publication of the sequence information for several thousand clones of the Kazusa TAC collection (Hirose et al., 2015), also available via the ABRC and RIKEN, and our Genome Browser tool (https://brcwebportal.cos.ncsu.edu/plant-riboprints/ArabidopsisJBrowser/) and out MATLAB application (https://github.com/Alonso-Stepanova-Lab/Recombineering-App) to identify the best TAC clone and set of primers to tag any given gene, should significantly improve the accessibility and use of recombineering in plants.

To extend the use of recombineering beyond *Arabidopsis*, we have also developed another set of recombineering cassettes and binary vectors for the efficient transfer of large fragments of DNA from a BAC to high-capacity transformation-ready vectors, such as derivatives of pYLTAC17. This opens the possibility of using recombineering in any transformable plant species for which a BAC library covering the whole genome has been at least end-sequenced. Previous work from the Dr. Csaba Koncz group has implemented the use of gap-repair cloning to transfer DNA from a BAC to binary vectors (Bitrian et al., 2011). Although this is a clever and relatively simple approach, it requires the cloning of different genomic DNA fragments in a binary vector for each GOI, limiting its convenience and scalability. Our cassette-exchange approach expands the ability to employ recombineering not only to other plant species, but also allows for scalability and the use in plant transformation of very large DNA fragments (over 75 kb) originally present in a BAC clone.

Finally, our antibiotic-based positive/negative selection cassettes (such as the *Universal tag-generator* cassette) provide a simple way to convert any existing tag into a recombineering-ready cassette. Thus, although our toolset comes with a collection of reporter tags ready to be used in gene expression analysis experiments, other types of specialized tags (such as those for the study of protein-protein interactions, protein-DNA, protein-RNA complexes, etc.) can be easily converted into recombineering cassettes using our tag-generator tool. This same tag-generator tool can also be utilized to make more sophisticated gene edits in the context of a BAC clone. In these types of experiments, the *tag generator* cassette is first inserted in the location near the point where the change needs to be introduced using the positive selection for ampicillin. The whole cassette can then be replaced by the sequence of choice by selecting against *RPSL* in the presence of streptomycin. The only limitation of the type of modification that can be made using this approach is the size of the DNA fragment used to replace the *Universal tag-generator* cassette due the inverse relationship between the size of a linear DNA fragment and its electroporation efficiency into *E. coli*. However, most applications only require the use of up to a few thousand base pairs as replacement DNA, and fragments of such sizes can be efficiently transformed into the recombineering *E. coli* strains. Thus, although the tag generator cassette is functionally equivalent to the classic *galK* cassette, it has the clear advantage of requiring simple LB medium and highly efficient antibiotic resistance instead of complicated and expensive minimum media and 2-deoxy galactose metabolic selection required for the use of the *galK*-based systems. In summary, the toolset and resources described in this work should make it possible for any molecular biology research laboratory, and even teaching laboratories equipped for basic bacterial growth and PCR amplification, to carry out large arrays of gene editing experiments by recombineering.

To further demonstrate the utility of the developed tools and resources, we implemented an experimental pipeline for tagging by recombineering 96 genes in parallel. Although high-throughput protocols have been previously developed for the generation of genome-wide translational fusions in *Drosophila* and *C. elegans*, we have opted for an intermediate throughput where individual clones after each transformation or recombination event are tested. We believe that the approach described here is better when a relatively small number of genes are being tagged, as it ensures that final constructs will be obtained for most, if not all, of the genes of interest. The testing steps in solid media, however, could be easily eliminated as has been done previously by others (Sarov et al., 2012; Sarov et al., 2016) to further increase the throughput of the procedure, although at the likely cost of a decrease in the percentage of genes finally tagged.

### Auxin biosynthesis

Auxin gradients play key roles in plant growth and development. In the past, the morphogenic auxin gradients have been mainly explained by a combined action of auxin transport and signaling/response (reviewed in (Vanneste and Friml, 2009)). Only in the last few years the contribution of local auxin production has been associated with the generation and maintenance of the morphogenic auxin maxima (Stepanova et al., 2008; Brumos et al., 2018; Zhao, 2018). Our present work characterizing the expression patterns of all the genes involved in IAA production through the indole-3-pyruvic acid (IPyA) pathway, the main route of auxin biosynthesis in *Arabidopsis* (Mashiguchi et al., 2011; Stepanova et al., 2011), sheds fresh light on the spatiotemporal patterns of auxin production by precisely defining the domains of activity of every *TAA1/TAR* and *YUC* gene in a limited set of tissues and developmental stages.

The establishment and maintenance of the shoot (SAM) and root apical meristems (RAM) is governed by auxin gradients generated by a joint action of local auxin biosynthesis and transport (Brumos et al., 2018; Wang and Jiao, 2018). Our observations indicate that in the SAM, auxin is locally synthesized by the tryptophan aminotransferases *TAA1* and *TAR2* and flavin monooxygenases *YUC1* and *YUC4*. In roots, *TAA1, YUC3, YUC7, YUC8* and *YUC9* are responsible for the production of IAA in the stem cell niche of the RAM. These observations are in agreement with recent single-cell RNA sequencing assays profiling the developmental landscape of *Arabidopsis* root (Zhang et al., 2019), where *YUC3*, *YUC8* and *YUC9* are included in the stem cell niche clusters.

In roots, ethylene triggers local auxin biosynthesis leading to an increase in auxin levels and the inhibition of root elongation (Stepanova et al., 2005; Ruzicka et al., 2007; Stepanova et al., 2007; Swarup et al., 2007; Stepanova et al., 2008; Brumos et al., 2018). Higher-order mutants of the *TAA1/TAR* and *YUC* gene families (Stepanova et al., 2008; Mashiguchi et al., 2011) display root-specific ethylene insensitive phenotypes. However, the specific genes involved in the local production responsible for the boost in auxin levels, particularly in the root elongation zone, have not been yet identified. Herein, we discovered that multiple genes of the IPyA pathway (*TAA1, TAR1, TAR2, YUC3, YUC5, YUC8*, and *YUC11*) are induced in roots by the treatment with the ethylene precursor ACC, with *TAA1, YUC3* and *YUC5* displaying a clear upregulation in the elongation zone. This observation suggests that auxin locally produced by these genes in the elongation zone may contribute to the arrest of root growth in the presence of ethylene. In addition, other ethylene-inducible *TAR*s and *YUC*s may also contribute to the ethylene-triggered auxin-mediated root growth inhibition, as auxin transport also plays an important role in the ethylene responses in roots via transcriptional induction of *AUX1, PIN1, PIN2*, and *PIN4* by ethylene (Ruzicka et al., 2007).

Our survey of auxin gene expression patterns in the recombineering fusions has unexpectedly uncovered the ACC-triggered induction of multiple auxin biosynthesis genes in the shoots of etiolated seedlings. As many as 10 of the 14 genes investigated, *TAR2, YUC2, YUC3, YUC4, YUC5, YUC6, YUC7, YUC8, YUC10*, and *YUC11*, are upregulated by ethylene in the hypocotyls and/or cotyledons, suggesting that a boost in auxin levels may contribute to the ethylene-induced shortening of hypocotyls and/or inhibition of cotyledon expansion (Vaseva et al., 2018). To date, the effect of ethylene on auxin biosynthesis has been extensively investigated only in roots (Stepanova et al., 2005; Ruzicka et al., 2007; Stepanova et al., 2007; Swarup et al., 2007; Stepanova et al., 2008; Brumos et al., 2018). Having the new recombineering reporter lines available for all major auxin biosynthetic pathway genes opens doors not only to the study of auxin production in seedlings, but also to the dissection of spatiotemporal patterns of local auxin biosynthesis in all organs and tissues under a myriad of different conditions, genotypes, and treatments. In fact, an inquiry into the spatial distribution of the expression of auxin biosynthetic genes in reproductive organs uncovered anthers, gynoecia, and developing ovules and seeds as the major sites of auxin biosynthesis. What is, perhaps, unexpected is that in the flowers, the vast majority of *YUC* gene activity (and, consequently, the expression of the auxin responsive reporter *DR5:GUS*) is concentrated almost exclusively in the male reproductive organs (in the anthers), whereas *TAA1* is predominantly active in the female organs (in the gynoecia). These observations suggest that some of the product of the TAA1/TAR-catalyzed biochemical reaction, IPyA, that serves as a substrate for YUCs to make auxin, IAA, may be transported within the flowers out of the gynoecia, e.g. to the anthers. With IPyA being a highly labile compound, at least *in vitro* (Tam and Normanly, 1998), determining if and how it moves within the plant may be challenging. Alternatively, *YUC* expression in the gynoecia may simply be below out detection limit, or the conversion of IPyA to IAA may not be the rate-limiting “bottleneck” step in every tissue that makes auxin. Nonetheless, some IPyA is likely made directly in young anthers, as *TAA1, TAR1* and *TAR2* all show some activity in those organs. The local anther-made IPyA, together with the pool of IPyA potentially transported from the gynoecia, can then be utilized by multiple anther-expressed *YUC*s to make auxin that contributes to pollen maturation, pre-anthesis filament elongation and anther dehiscence (Cecchetti et al., 2008). Our prior work (Brumos et al., 2018) indicates that the spatiotemporal misregulation of *TAA1* expression in developing flowers, that is expected to shift the domains of local IPyA and, hence, auxin production, results in flower infertility, highlighting the importance of the specific patterns of auxin gene activity for proper flower development. With the new recombineering resources at hand, we can now start dissecting the roles of individual *TAA1/TAR* and *YUC* family members in the development of flowers and other organs and tissues in *Arabidopsis*.

Several of the new translational reporters generated in this study by recombineering, specifically those for *TAA1*, *YUC1*, *YUC2, YUC3, YUC4, YUC5* and *YUC7*, behave differently from the previously published transcriptional reporters for the same genes (Cheng et al., 2006; Yamada et al., 2009; Lee et al., 2012; Chen et al., 2014; Challa et al., 2016; Kasahara, 2016; Brumos et al., 2018; Xu et al., 2018). For example, in primary roots, *TAA1* transcriptional fusions are mainly expressed in the stele (Yamada et al., 2009; Brumos et al., 2018), whereas the recombineering construct is active in the quiescent center (QC) and pro-vasculature (this work and (Stepanova et al., 2008; Brumos et al., 2018)). *YUC3* promoter fusion is the strongest in the elongation zone of the primary root (Chen et al., 2014), but the recombineering construct is predominantly detected in the QC, as well as in the columella initials and the root cap (this work). For *YUC5*, prominent QC expression is seen for transcriptional fusions (Challa et al., 2016), but not for translational fusions generated by recombineering (this work), yet both constructs are active along the edges of the cotyledons. For *YUC7*, a transcriptional fusion is mildly active in the proximal regions of the root but is not detectable in the root meristem (Lee et al., 2012), whereas the recombineering construct for this gene is highly active in the QC and the root cap (this work). Analogous discrepancies are seen in the reproductive organs. For example, in mature flowers, *YUC1* is detectable in the flower abscission zones only with a recombineering translational fusion (this work), but not with a transcriptional reporter (Cheng et al., 2006). *YUC2* promoter fusion shows expression in young flowers, specifically in the valves of gynoecia, the pedicels, flower organ abscission zones, and petals, and no detectable expression in mature flowers (Cheng et al., 2006), whereas a recombineering translational fusion construct for this gene is active in the anthers of young flowers and in the abscission zones of petals and sepals in mature flowers (this work). For *YUC4*, both transcriptional and translational fusions are expressed in the female reproductive structures, specifically in the stigmas, but in the male reproductive structures, the transcriptional reporters are active only in the distal tips of the anthers (Cheng et al., 2006), whereas the recombineering-generated translational fusions have more ubiquitous, uniform activity throughout the entire anthers (this work). In addition, transcriptional reporters for *YUC4* are detected in young flower buds at the base of floral organs (Cheng et al., 2006), but this domain of activity is not readily observed in the recombineering-generated translational reporter fusions (this work).

The differences in the expression patterns and levels of the transcriptional and translational reporters are likely due to the lack of some key regulatory elements in the promoter-only fusions that are captured in the recombineering reporters made in this work, as the latter constructs include much larger upstream (10kb) and downstream (5kb) regions of the genes and possess the full coding regions with all of the introns. In the recombineering constructs, the presence of introns can provide a diverse population of mRNAs due to alternative splicing, as may be the case for *YUC4* (Kriechbaumer et al., 2012). Differences in the length, content and structure of the transcripts can lead to differences in RNA stability and localization, resulting in variable expression levels and patterns (Kriechbaumer et al., 2012). It is, however, not uncommon to see discrepancies in the expression patterns of transcriptional versus translational fusions even for the constructs that harbor identical promoter fragments. For example, the activities of *YUC1* and *YUC4* reporters made by classical cloning approaches (Xu et al., 2018) differ for the transcriptional versus translational constructs, with translational reporters showing less activity specifically in young flowers than their respective transcriptional fusions. The differences in expression can be explained by the presence of negative transcriptional regulatory elements in the introns of these genes harbored in the translational fusions, as well as by the mobility and/or instability of fusion RNAs or proteins. For example, protein turnover plays a major role in the expression of auxin co-receptor proteins Aux/IAAs, with translational fusions, unlike transcriptional reporters, for these genes being hard to detect due to rapid Aux/IAA protein degradation in the presence of auxin (Tiwari et al., 2001; Zhou et al., 2011). Regardless of the molecular underpinnings of the expression pattern differences between previously published classical and newly generated recombineering reporters, the latter constructs harbor most if not all of the regulatory elements of the native gene and thus should be considered the gold standard in gene functional studies.

## METHODS

### General recombineering procedures

Recombineering experiments were carried out as described in (Alonso and Stepanova, 2015). In brief, SW105 cells carrying the TAC or BAC of interest were grown overnight at 32°C in LB supplemented with the antibiotic needed to select for the corresponding BAC or TAC. Overnight culture (1 mL) was used to inoculate 50 mL of LB plus antibiotic in a 250 mL flask and grown at 32°C for 2-3 h with constant shacking. The lambda-red recombineering system was activated by incubating the cells in a water bath at 42°C and constant shaking for 15 min. Cells were immediately cooled down in water-ice bath and electrocompetent cells were then prepared (Alonso and Stepanova, 2015). Cells were electroporated with the PCR-amplified recombineering cassette and allowed to recover in LB for 1 h at 32°C, and then plated in an LB plate with the corresponding selection. After a two-day incubation at 32°C, the presence of the recombination event in the primary transformants was confirmed by colony PCR using a gene-specific primer and a primer specific for the inserted cassette. Primer sequences are provided in Tables S1-S4. FLP reaction was carried out by growing the SW105 cells harboring the TAC or BAC clone with the desired recombineering cassette already inserted in the location of interest overnight at 32°C under constant shacking in LB media supplemented with the necessary antibiotics to select for the BAC or TAC. Fresh LB media (1 mL) with the antibiotic necessary to select for the BAC or TAC backbone was inoculated with 50 µL of overnight culture and 10 µL of 10% (w/v) L-arabinose. The cells were then grown for 3 hours at 32°C with constant shacking. A sterile toothpick was dipped in the culture and used to streak a few cells into a fresh LB plate supplemented with the antibiotic necessary to select for the BAC or TAC backbone with the goal of obtaining isolated colonies. Colonies were then tested by colony PCR to confirm the elimination of the *FRT*-flanked DNA sequences. To ensure that the modification in the GOI is correct, the test PCR product was sequenced using the corresponding test oligos.

Commercial DNA synthesis services (IDT) were used to obtain the following sequences: *Universal GUS-FRT-Amp-RFP*, *Universal RPSL-Amp* and *Universal tag-generator* cassettes, as well as the *Universal AraYpet*, and the *Universal 3xAraYpet* fluorescent protein genes. Sequences of these cassettes are provided in Supplemental Table S5.

### High-throughput recombineering and trimming of the *3xYPET* and *GUS* cassettes

The basic recombineering procedures were followed during the parallel processing of 96 constructs with the following modifications. The 96 DH10B strains carrying the genes of interest were inoculated in 96 1-mL LB kanamycin cultures in a 96-deep-well plate and grown overnight. TAC DNA was extracted by regular alkaline lysis (Alonso and Stepanova, 2014) using 12 strips of eight 1-mL tubes. In parallel, freshly prepared SW105 electrocompetent cells (Alonso and Stepanova, 2015) were aliquoted into the 96 electroporation wells of a 96-well electroporation plate (BTX electroporation Systems). 40 µL of competent cells and 3 µL of DNA were mixed in the cuvette and electroporated as previously described (Alonso and Stepanova, 2015). Cells were transferred to a 96-deep-well plate and incubated at 32°C with shaking for 1-2 hours to recover. Cells were collected by centrifugation and plated on LB kanamycin plates. After confirming by PCR the presence of the TAC clones using the testing primers (see Tables S1-S4), glycerol stocks for the 96 SW105 strains were generated. Using these stocks, 96 cultures were grown overnight in a 96-deep-well plate. Eight sets of 12 strains were pooled together to inoculate eight 250-mL flasks with LB kanamycin, grown for additional 3 hours, heat shocked at 42°C, and then used to prepare electrocompetent cells as previously described (Alonso and Stepanova, 2015). In parallel, 96 amplicons corresponding to the desired recombineering cassette (*Universal Venus-FRT-galK-FRT*, *Universal AraYpet-FRT-Amp-FRT* or *Universal 3xAraYpet-FRT-Amp-FRT*) were obtained using DNA template for the cassette and the corresponding recombineering primers (Tables S1 to S4). PCR fragments were purified by chloroform extraction and ethanol precipitation. PCR DNA was resuspended in 20 ul of water and 3 ul were used for electroporation in the 96-well electroporation cuvette as described above. Recovery, plating, and testing was also done as described above except that LB kanamycin and ampicillin plates were used for the selection. Test PCR products were sequenced to confirm the integrity and fidelity of the recombination events. For the trimming, the *Tet^R^* gene was amplified from the *FRT2-Tet-FRT2* trimming cassette to generate 96 trimming amplicons using the primers replaLB-tet Universal and one of the 96 Gene-DelRight primers (Table S4). The 96-well format recombination procedure was done as described above but selecting the recombination events in LB plate supplemented with kanamycin and tetracycline. The insertions were confirmed using the LBtest and the corresponding TestDelRigh primer (Table S4). A second round of trimming was carried out using 96 amplicons obtained by amplifying the *Amp^R^* gene from the *FRT5-Amp-FRT5* trimming cassette with the primers replaRB-amp Universal and one of the 96 Gene-DelLeft recombineering primers. After confirming the trimming by PCR using the primers testRB and the corresponding TestDelLeft oligo, the second antibiotic resistance gene, *aadA*, an *aminoglycoside 3’-adenylyltransferase* gene that confers spectinomycin and streptomycin resistance in both in *E. coli* and *Agrobacterium*, was introduced in the trimmed constructs by recombineering using an amplicon obtained by amplifying the *Kan-Spec cassette* using the primers Spect-Kan test F and Spect-Kan test R (Table S4). Plasmid DNAs for the 96 strains obtained were prepared by alkaline lysis using 12 strips of eight 1-mL tubes and electroporated into electrocompetent *Agrobacterium* using the same 96-well procedure as described above. *Agrobacterium* selection was done using LB pates supplemented with kanamycin and spectinomycin.

### Generation of the Universal Venus-FRT-galK-FRT cassette

The *Universal AraYpet* was utilized as a template with the primers PEO1F and PEO1R. This amplicon was inserted in the JAtY clone JAtY68N23 using the classical *galK* system as described (Zhou et al., 2011) to generate the *Universal AraYpet* cassette. The *Venus-FRT-galK-FRT* sequences from the pBalu6 were amplified using the primers VenusPeo1F and VenusPeo1R and this PCR product was employed to replace the *Ypet* sequences in the *Universal AraYpet* cassette by recombineering, as described (Zhou et al., 2011).

### Generation of the Universal mCherry-FRT-galK-FRT cassette

A strategy similar to that used to generate the *Universal Venus-FRT-Galk-FRT* was utilized to produce the *Universal mCherry-FRT-galK-FRT* cassette, but in this case the primers CherryPeo1F and VenusPeo1R were employed to amplify the *mCherry FRT-galK-FRT* sequences from pBalu8. This PCR product then served to replace the *Ypet* sequences in the *Universal AraYpet cassette* by recombineering as described (Zhou et al., 2011).

### Generation of the Universal AraYpet-FRT-Amp-FRT cassette

The ampicillin resistance gene, *Amp^R^*, and the corresponding promoter were amplified from pBluescript with the primers PEO1FRTAmpF and PEO1FRTAmpR. This PCR product was then used in a recombineering reaction to insert the *FRT-Amp-FRT* sequences in the *Universal AraYpet* cassette to generate the *Universal AraYpet-FRT-Amp-FRT* cassette.

### Generation of the Universal AraYpet-FRT-TetA-FRT cassette

The tetracycline resistance gene, *Tet^R^*, and the corresponding promoter sequences were amplified from the genomic DNA of the recombineering strain of *E. coli* SW102 with the primers PEO1FRTtetAF and PEO1FRTtetRAR. This PCR product was then employed in a recombineering reaction to insert the *FRT-Tet-FRT* sequences in the *Universal AraYpet* cassette to generate the *Universal AraYpet-FRT-TetA-FRT* cassette.

### Generation of the Universal *3xAraYpet-FRT-Amp-FRT* cassette

The *Universal 3xAraYpet* sequence was commercially synthesized by IDT and utilized as a template for a PCR reaction with the primers IAA5F and IAA5R. The obtained amplicon was inserted in the JAtY61G08 clone using the classical *galK* recombineering approach as described (Zhou et al., 2011) to generate the *Universal 3xAraYpet* cassette. The *FRT-Amp-FRT* sequences from the *Universal AraYpet-FRT-Amp-FRT* cassette were amplified with the primers 3YpetFAFF1 and 3YpetFAFR1 and inserted in the *Universal 3xAraYpet* cassette to create the *Universal 3xAraYpet-FRT-Amp-FRT* cassette.

### Generation of the *Universal-RFP-FRT-Amp-FRT* cassette

The *Universal tag-generator* cassette was commercially synthesized by IDT and then amplified with the primers PCL5_STOP_5UA and 3UA_PCL3. The resulting amplicon was inserted in the tomato BAC clone HBa0079M15 by recombineering (Zhou et al., 2011). The *RFP* DNA was amplified from the pUBC-RFP-DEST vector (Grefen et al., 2010) with the primers PCL5_5UA_RFP-f and UR_RFP-r. The resulting product was used in a recombineering reaction to replace the *RPSL* in the BAC clone containing the *Universal tag-generator* cassette.

### Generation of the pDONR221-FRT2-SacB-FRT5 and pGWB1-FRT2-SacB-FRT5 vectors

The *SacB* gene was amplified from the BiBAC2 vector (Hamilton, 1997) with the primers FRT2SLongNew and FRT5Long. The PCR product was then cloned in the pDONR221 to create the pDONR221-FRT2-SacB-FRT5 vector. Spacer sequences for the orthogonal *FRT*s were obtained from a published source (Schlake and Bode, 1994). The pGWB1-FRT2-SacB-FRT5 vector was generated by transferring the *FRT2-SacB-FRT5* sequences from the pDONR221-FRT2-SacB-FRT5 to the pGWB1 binary vector using a Gateway LR reaction.

### Generation of the *Kan-Spec* cassette

The *aadA* sequences were amplified from the pTF101 vector with the primers SpectFKan and SpectRKan and inserted in the JAtY63D14 clone to generate the *Kan-Spec* cassette template. Primers Spect-Kan test F and Spect-Kan test R can be used to amplify the *Kan-Spec* cassette from the *Kan-Spec* cassette template.

### Generation of the pYLTAC17-FRT2-SacB-FRT5-Spec vector

The *RPSL-Amp* sequences were amplified from the *Universal RPSL-Amp* cassette with the primers replaRBAmpRPSL and replaLBTetRPSL. The resulting PCR product was used to replace all of the *Arabidopsis* genomic and the *SacB* sequences in the JAtY56F21 clone by recombineering (Zhou et al., 2011) producing the pYLTAC17-RPSL-Amp vector. Next, the *FRT2-SacB-FRT2* cassette was amplified from the pDONR221-FRT2-SacB-FRT5 vector with the primers replaRBSacBM13F and replaLBSacBM13R and utilized in a recombineering reaction to replace the *RPSL-Amp* sequence in the pYLTAC17-RPSL-Amp to generate the pYLTAC17-FRT2-SacB-FRT5 vector. Next, the bacterial *aadA* (*aminoglycoside-3’-adenyltransferase*) gene to confer spectinomycin and streptomycin resistance was amplified from *Spec-Kan* cassette by PCR with primers Spec-Kan test F and Spec-Kan test R and integrated into the pYLTAC17-FRT2-SacB-FRT5 vector by recombineering to generate the final pYLTAC17-FRT2-SacB-FRT5-Spec vector.

### Generation of the pYLTAC17-FRT2-SacB-FRT5-Spec-Kan vector

The *RPSL-Amp* sequences were amplified from the *Universal RPSL-Amp* cassette with the primers pYLTAC-RPSL-Amp-f2 and pYLTAC-RPSL-Amp-r2. The resulting PCR product was used to replace the Act1 5’-Bar-Nos 3’ cassette in pYLTAC17-FRT2-SacB-FRT5 to generate the pYLTAC17-FRT2-SacB-FRT5-RPSL-Amp. Next, the *Kan* resistance cassette for plant selection was amplified from pGWB1 by PCR with primers pYLTAC-Kan-f2 and pYLTAC-Kan-r2, and the resulting PCR product was utilized to replace the *RPSL-Amp* sequence in pYLTAC17-FRT2-SacB-FRT5-RPSL-Amp to generate pYLTAC17-FRT2-SacB-FRT5-Kan. Finally, the bacterial *aadA* gene to confer spectinomycin and streptomycin resistance was amplified from the *Spec-Kan* cassette by PCR with primers Spec-Kan-testF and Spec-Kan-testR and integrated into the pYLTAC17-FRT2-SacB-FRT5-Kan vector by recombineering to generate the final pYLTAC17-FRT2-SacB-FRT5-Spec-Kan vector.

### Generation of the *FRT2-Tet-FRT2* trimming cassette

The *tetA* resistance gene was amplified from the *Universal AraYpet-FRT-TetA-FRT* cassette with the primers FRT2-Tet-F (JAtY Universal) and FRT2-Tet-R (EIN3del) and inserted in the TAC clone JAtY63D14 by recombineering.

### Generation of the *FRT5-Amp-FRT5* trimming cassette

The ampicillin resistance gene, *Amp^R^*, was amplified with the primers FRT5-Amp-R (BeloBAC11Right universal) and FRT5-Amp-PIN5delR. The resulting PCR product was used as a template in a second PCR reaction with the primers FRT5-Amp-EIN3delR and FRT5-Amp-R (JAtY universal) and the resulting PCR product was employed in a recombineering reaction resulting in the insertion of the *FRT5-Amp-FRT5* cassette in the TAC clone JAtY63D14.

### Generation of the Universal galK-FRT-Amp-FRT cassette

The *galK* sequence was amplified from the *Universal Venus-FRT-galK-FRT* cassette with the primers UnigalK F and UniGalk R. The resulting PCR product was utilized in a recombineering reaction to replace the *AraYpet* sequences from the *Universal AraYpet-FRT-Amp-FRT* cassette resulting in the *Universal galK-FRT-Amp-FRT* cassette.

### Generation of the Universal GFP-FRT-Amp-FRT, Universal mCherry-FRT-AmpFRT, and Universal 3xMYC-FRT-Amp-FRT cassettes

The sequences of the *GFP*, *mCherry* and *3xMYC* were amplified with the primer pairs UniGFP F/UniGFP R, mCherryAmpF/mCherryAmpR and Uni3XMYC F/Uni3XMYC R, respectively. Each of the PCR products was used in an independent recombineering experiment to replace the *galK* sequence of the *Universal Galk-FRT-Amp-FRT* cassette by the sequences of each of these new tags.

### Generation of the Universal AraYpet-3xMYC-FRT-Amp-FRT

The *3xMYC-FRT-Amp-FRT* sequence was amplified from the *Universal 3xMYC-FRT-Amp-FRT* cassette with the primers Ypet-3xMYC and Uni3xMYC R. The corresponding PCR product was inserted immediately after the *Ypet* sequence to generate the *Universal AraYpet-3xMYC-FRT-Amp-FRT* cassette.

### Examination of the *in vivo* efficiency of an exchange cassette reaction

To evaluate the efficiency of transferring large DNA fragments from a BAC clone to two different binary vectors, pGWB1-FRT5-SaB-FRT5 and pYLTAC17-FRT2-SacB-FRT5-Spec, the *YUC9* gene (*At1g04180*) was tagged with *GUS* in the C-terminus by a recombineering reaction in which the *Universal AraGus-FRT-Amp-FRT* cassette was amplified with the primers InsertGUS-Amp-f and InsertGUS-Amp-r. To generate DNA sequences (that contain *YUC9-GUS*) of different sizes flanked by the *FRT2* and *FRT5* sites, the *FRT2-Tet-FRT2 trimming* and *FRT5-Amp-FRT5 trimming* cassettes were inserted 10, 25, or 57 kb upstream and 5, 11 and 20 kb downstream of the *YUC9* gene, respectively, by recombineering, where the *FRT2-Tet-FRT2 trimming* cassette was amplified with the primer pairs Up-10kb_FRT2_f/ Up-10kb_FRT2_r, Up-25kb_FRT2_f/ Up-25kb_FRT2_r and Up-57kb_FRT2_f/ Up-57kb_FRT2_r, and the *FRT5-Amp-FRT5 trimming* cassette was amplified with the primer pairs Down-5kb_FRT5_f/ Down-5kb_FRT5_r, Down-11kb_FRT5_f/ Down-11kb_FRT5_r and Down-20kb_FRT5_f/ Down-20kb_FRT5_r. After inserting the corresponding PCR fragments in the BAC clone and removing the antibiotic resistance genes in a FLP reaction, three clones containing the *YUC9* gene tagged with *GUS* in the C-terminus and flanked by 10kb upstream and 5 kb downstream, 25 kb upstream and 11 kb downstream, or 57 kb upstream and 20 kb downstream with the *FRT2* and *FRT5* sites, respectively, were generated. The *in vivo* cassette exchange reaction was carried out by electroporating *E. coli* SW105 competent cells carrying one of the three *YUC9-GUS* constructs described above and grown in the presence of 0.1% (w/v) L-arabinose for the three hours prior to starting the process of preparing the cultures for electroporation. After the electroporation with either the pGWB1-FRT5-SaB-FRT5 and pYLTAC17-FRT2-SacB-FRT5-Spec vectors, the cells were allowed to recover for additional 3 hours at 32°C in LB media supplemented with 0.1% (w/v) L-arabinose. Clones containing the binary vectors carrying the *YUC9-GUS* genomic sequences delimited by the *FRT2* and *FRT5* sites were selected in LB plates supplemented with 10% (w/v) sucrose (to select against the unmodified binary vectors) and either kanamycin (50 mg/mL) and hygromycin (200mg/mL) to select for the pGWB1-based plasmids or kanamycin (50 mg/mL) and spectinomycin (50 mg/mL) to select for the pYLTAC17-derived vectors, respectively.

### Plant transformation and fluorescence analysis

Transformation of the *Agrobacterium tumefaciens* strain UIA143 pMP90 (Hamilton, 1997; Hamilton, 1997) with the tagged constructs of interest was performed by electroporation as described (Alonso and Stepanova, 2015). The resulting colonies were re-streaked on LB plates supplemented with the appropriate antibiotic and single colonies were tested by PCR with gene-specific primers to confirm presence of the construct. Fresh colonies were inoculated into 5 mL of liquid LB plus kanamycin, grown with shaking overnight at 28°C, and the resulting saturated cultures were split into two and plated onto two 150mm LB kanamycin plate. After two nights at 30°C, the cells were scraped off with a spatula and resuspended in 100 mL of liquid transformation solution (1xMS (pH6.0) 1% (w/v) glucose spiked with 200 µL/L Silvett-77). Wild-type *Arabidopsis* (Col-0) grown in soil under 16 h light/ 8 h dark cycle until inflorescences are about 15 cm long were transformed with the resulting cultures using flower dip method (Clough and Bent, 1998). Plants were allowed to recover under a plastic dome for 24-48 h, and then grown to maturity. T1 plants were selected in 20 µg/mL phosphinothricin in AT plates (1xMS, 1% (w/v) sucrose, pH6.0 with KOH, 7% (w/v) bactoagar) and propagated in soil (50:50 mix of Sun Gro gemination and propagation mixes) under 16 h light/ 8 h dark cycle. T3 plants homozygous for the constructs were confirmed by genotyping with a combination of tag-specific (*Ypet* or *GUS*) and gene-specific primers.

Fluorescence analysis of *Ypet*- and *3xYpet*-tagged lines was performed using a Zeiss Axioplan microscope in the T1, T2, and/or T3 generations, focusing on the expression patterns in three-day-old etiolated seedlings. T3 lines homozygous for the tagged constructs listed in Supplemental Tables S1 to S3 were donated to the Arabidopsis Biological Resource Center.

### GUS staining and optical clearing of plant tissues

Seeds of homozygous T3 and/or T4 *GUS*-tagged lines were sterilized with 50% (v/v) commercial bleach spiked with Triton to break seed clumps, washed five or more times with sterile water to remove bleach, resuspended in melted and precooled sterile 0.7% (w/v) low-melting point agarose, and plated on plain AT plates or plates supplemented with 10 µM ACC, 10 µM NPA, 10 µM ACC plus 10 µM NPA, or 50 nM NAA. After three days at 4°C to equalize germination, seedings were exposed to light for 1-2 hours at room temperature to restart the clock and germinated at 22°C for three days in the dark. Seedlings were fixed in cold 90% acetone and immediately stained for GUS overnight as described (Stepanova et al., 2005). For flower and inflorescence analysis, transgenic T3 and T4 lines homozygous for the transgenes were grown in soil under 16 h light/ 8 h dark cycle. Tips of inflorescences (∼3 cm) were excised with small scissors, fixed in cold 90% (v/v) acetone, stored overnight at −20°C to help remove chlorophyll, stained for GUS as described (Stepanova et al., 2005), and then stored in 70% EtOH for several additional days to remove residual chlorophyll prior to imaging. Etiolated 3-day-old seedlings were fixed in 90% (v/v) acetone and optically cleared using freshly prepared ClearSee solution (Kurihara et al., 2015) for at least 7 days. Images of inflorescences mounted on AT plates were taken with Q Capture software on a 5.0 RTV digital camera (Q Imaging, Surrey, BC, 904 Canada) under a Leica MZ12.5 stereomicroscope. To examine roots, hypocotyls, and flowers, samples were mounted on glass slides and imaged with the same camera and software on a Zeiss AxioSkop2 Plus microscope with Nomarski optics.

### Practical guide for standard recombineering applications

To make recombineering more accessible to plant researchers, we provide step-by-step instructions on how to implement this method using the resources and tools reported in this study.

#### (A) Perform a standard gene tagging experiment with one of the Universal recombineering cassettes of the collection

In order to insert a tag from the collection using our Universal recombineering cassettes, first a TAC (using our Genome Browser https://brcwebportal.cos.ncsu.edu/plant-riboprints/ArabidopsisJBrowser/ or MATLAB application https://github.com/Alonso-Stepanova-Lab/Recombineering-App for Arabidopsis) or BAC clone containing a GOI needs to be identified. (2) Next, the TAC/BAC DNA is isolated and transferred via electroporation to the SW105 recombineering strain. (3) Recombineering and testing primers for the GOI are designed either using our Genome Browser or by generating the following forward and reverse primers: Recombineering-F: 5’-40 nt identical to the sequence immediately upstream of the desired insertion point for the tag followed by the sequence - GGAGGTGGAGGTGGAGCT-3’; Recombineering-R: 5’- reverse complement of the 40 nt immediately downstream of the desired insertion point followed by the sequence - GGCCCCAGCGGCCGCAGCAG-3’. (4) The next step is to generate a recombineering amplicon using these primers, any of the recombineering cassettes with the tag of interest as a template, and a proofreading polymerase. (5) The amplicon is then inserted into the desired location by recombineering as described above in the *General recombineering procedures* methods section. (6) To test the resulting colonies, test primers flanking the insertion site and regular Taq polymerase are employed for colony PCRs on the recombineering products. (7) Once a desired clone is identified, the antibiotic selection sequences in the tag are removed using an *in vivo* FLP reaction, as described in the *General recombineering procedures* methods section above. (8) Finally, the construct is verified by re-sequencing of the integrated DNA tag and the genomic DNA-tag junction sites using the test primers.

#### (B) Perform a trimming experiment of a TAC or BAC clone using the FRT2-tet-FRT2 and FRT5-Amp-FRT5 cassettes

(1, 2) The first two steps of the procedure as the same as above in section (A). (3) Then, the *FRT2-Tet-FRT2* cassette is amplified from the template using the primers FRT2-Tet-FRT2-F and FRT2-Tet-FRT2-R. The forward FRT2-Tet-FRT2-F primer is comprised of: 5’- 40 nt identical to the 40 nucleotides upstream of the sequence to be deleted at the 5’-end of the clone followed by the sequence AACGAATGCTAGTCTAGCTG-3’. For example, when using the JAtY or KAZUSA TAC, we use the primer NewreplaRBFRT2-Tet 5’-TATATTGCTCTAATAAATTTTTGGCGCGCCGGCCAATTAGGCCCGGGCGG-TTCAAATATGTATCCGCTCATG-3’. Similarly, the reverse FRT2-Tet-FRT2-R primer consists of: 5’- 40 nt with the reverse complement sequence just downstream of the sequence to be deleted at the 3’-end of the TAC or BAC clone followed by the sequenceTTACCAATGCTTAATCAGTG-3’. (4) In parallel, the *FRT5-Amp-FRT5* cassette is amplified using the primers FRT5-Amp-FRT5F and FRT5-Amp-FRT5R. These consist of: 5’- 40 nt identical to the 40 nucleotides upstream of the sequence to be deleted at the 3’-end of the TAC/BAC clone followed by the sequence AACGAATGCTAGTCTAGCTG-3’ for the forward primer and 5’- 40 nt with reverse complement to the 40 nucleotides downstream of the sequence to be deleted at the 3’-end of the TAC/BAC clone followed by the sequence-TTAGTTGACTGTCAGCTGTC-3’ for the reverse primer. When using the JAtY or KAZUSA libraries, the primer NewreplaLBFRT5-Amp 5’- TTAGTTGACTGTCAGCTGTCCTTGCTCCAGGATGCTGTTTTTGACAACGG-TTAGTTGACTGTCAGCTGTC-3’ is used as a reverse primer. After generating these amplicons, steps (5) to (8) from section (A) are followed to trim and confirm the desired deletions.

#### (C) Generate a recombineering cassette for a new tag using the Universal tag-generator cassette

To generate a new tag, the Universal tag-generator cassette is provided as a ready-to-use SW105 strain carrying the tomato BAC clone HBa0079M15 harboring the Universal tag-generator cassette (Figure 2A). The following steps are required to generate any new tag. (1) The tag of interest (e.g., LUCIFERASE) is amplified from a DNA template with a proofreading polymerase and the primers tag-generatorF and tag-generatorR. These consist of the sequence 5’-TAAAAAGGGTTCTCGTTGCTAAGGAGGTGGAGGTGGAGCT-followed by the first 20 nucleotides of the tag of interest-3’ (it is important that the sequence of the tag starts with the first nucleotide of the first codon of the tag) for the forward primer and 5’- GAAAGTATAGGAACTTCCCACCTGCAGCTCCACCTGCAGC- followed by the reverse complement of the last 20 nucleotides before the stop codon of the tag of interest-3’ for the reverse primer. (2) To replace the RPSL marker in the Universal tag-generator cassette by the Tag-generator amplicon from the previous step, a standard recombineering protocol is employed as described in the General recombineering procedures methods section, except that the positive recombinant colonies are selected in LB media supplemented with streptomycin to select against the RPSL gene. (3) The sequence integrity of the new tag, as well as that of the recombination sites, is confirmed by sequencing a PCR product using primers flanking the new tag.

#### (D) Perform a sequence replacement/deletion using the Universal tag-generator or the Universal RPSL-Amp cassettes

(1, 2) The first two steps of the procedure are the same as above in section (A). (3) The *Universal tag-generator* or *Universal RPSL-Amp* amplicons are generated using a replacementF primer with the sequence 5’-40 nt upstream of the sequence to be modified followed by GGAGGTGGAGGTGGAGCT-3’ and a replacementR primer with the sequence 5’- 40 nt reverse complementary to the 40 nt immediately downstream of the sequence to be modified GGCCCCAGCGGCCGCAGCAG -3’. (4) The sequence to be modified in the TAC/BAC clone is replaced with the amplicon from step (3) using the standard recombineering protocol in the *General recombineering procedures* methods section selecting for ampicillin-resistant colonies. (5) A DNA fragment is commercially synthesized that contains the sequence with the desired modifications/deletions flanked by at least 40 nt (preferably, 100 to 200 nt) of the sequences homologous to the sequences flanking the inserted amplicon. (6) The replacement sequences from the previous step are employed to replace the amplicon inserted in step (4) using the same recombineering procedures as described above in the section *Generate a recombineering cassette for a new tag using the Universal tag-generator cassette* for the replacement of *RPSL* by the tag of interest.

## Acknowledgements

We are grateful to Jeonga Yun, Clara Alonso-Stepanova, Cierra Clark, Katherine Hobbet, and Emily Nelson for technical assistance, to Dr. Robert Franks for making his equipment available for this research, to Dr. Hobert for providing the *Venus-FRT-galK-FRT* cassette, to Dr. Ian Bancroft for providing the JAtY TAC collection, and to Dr. Donald Court and the National Cancer Institute for providing the SW105 strain. This work was supported by the NSF grants DBI 0820755 and MCB0519869 to JMA, MCB0923727 and IOS1444561 to JMA and ANS, and a Programa de Movilidad fellowship PRX14/from the Spanish Ministry of Science, Culture and Sports to MAPA.

## AUTHORS CONTRIBUTIONS

JMA, ANS, JB, CZ and MAPA designed the experiments and performed research. JMA, ANS, JB and CZ wrote the manuscript. YG, DS, and APP assisted in research.

## REFERENCES

Alonso, J.M. and Stepanova, A.N. (2014). Arabidopsis transformation with large bacterial artificial chromosomes. Methods Mol. Biol. 1062, 271–283, doi/10.1007/978-1-62703-580-4_15 [doi].

Alonso, J.M. and Stepanova, A.N. (2015). A recombineering-based gene tagging system for Arabidopsis. Methods Mol. Biol. 1227, 233–243, doi/10.1007/978-1-4939-1652-8_11 [doi].

Alvarez-Buylla, E.R., Benitez, M., Corvera-Poire, A., Chaos Cador, A., de Folter, S., Gamboa de Buen, A., Garay-Arroyo, A., Garcia-Ponce, B., Jaimes-Miranda, F., Perez-Ruiz, R.V., Pineyro-Nelson, A., and Sanchez-Corrales, Y.E. (2010). Flower development. Arabidopsis Book 8, e0127, doi/10.1199/tab.0127 [doi].

Band, L.R., Wells, D.M., Fozard, J.A., Ghetiu, T., French, A.P., Pound, M.P., Wilson, M.H., Yu, L., Li, W., Hijazi, H.I., Oh, J., Pearce, S.P., Perez-Amador, M.A., Yun, J., Kramer, E., Alonso, J.M., Godin, C., Vernoux, T., Hodgman, T.C., Pridmore, T.P., Swarup, R., King, J.R., and Bennett, M.J. (2014). Systems analysis of auxin transport in the Arabidopsis root apex. Plant Cell 26, 862–875, doi/10.1105/tpc.113.119495 [doi].

Begemann, M.B., Gray, B.N., January, E., Gordon, G.C., He, Y., Liu, H., Wu, X., Brutnell, T.P., Mockler, T.C., and Oufattole, M. (2017). Precise insertion and guided editing of higher plant genomes using Cpf1 CRISPR nucleases. Sci. Rep. 7, 11606-017-11760-6, doi/10.1038/s41598-017-11760-6 [doi].

Bhosale, R., Giri, J., Pandey, B.K., Giehl, R.F.H., Hartmann, A., Traini, R., Truskina, J., Leftley, N., Hanlon, M., Swarup, K., Rashed, A., Voss, U., Alonso, J., Stepanova, A., Yun, J., Ljung, K., Brown, K.M., Lynch, J.P., Dolan, L., Vernoux, T., Bishopp, A., Wells, D., von Wiren, N., Bennett, M.J., and Swarup, R. (2018). A mechanistic framework for auxin dependent Arabidopsis root hair elongation to low external phosphate. Nat. Commun. 9, 1409-018-03851-3, doi/10.1038/s41467-018-03851-3 [doi].

Bitrian, M., Roodbarkelari, F., Horvath, M., and Koncz, C. (2011). BAC-recombineering for studying plant gene regulation: developmental control and cellular localization of SnRK1 kinase subunits. Plant J. 65, 829–842, doi/10.1111/j.1365-313X.2010.04462.x; 10.1111/j.1365-313X.2010.04462.x.

Brumos, J., Robles, L.M., Yun, J., Vu, T.C., Jackson, S., Alonso, J.M., and Stepanova, A.N. (2018). Local Auxin Biosynthesis Is a Key Regulator of Plant Development. Dev. Cell. 47, 306–318.e5, doi/S1534-5807(18)30782-2 [pii].

Budiman, M.A., Mao, L., Wood, T.C., and Wing, R.A. (2000). A deep-coverage tomato BAC library and prospects toward development of an STC framework for genome sequencing. Genome Res. 10, 129–136.

Cecchetti, V., Altamura, M.M., Falasca, G., Costantino, P., and Cardarelli, M. (2008). Auxin regulates Arabidopsis anther dehiscence, pollen maturation, and filament elongation. Plant Cell 20, 1760–1774, doi/10.1105/tpc.107.057570 [doi].

Cermak, T., Baltes, N.J., Cegan, R., Zhang, Y., and Voytas, D.F. (2015). High-frequency, precise modification of the tomato genome. Genome Biol. 16, 232-015-0796-9, doi/10.1186/s13059-015-0796-9 [doi].

Challa, K.R., Aggarwal, P., and Nath, U. (2016). Activation of YUCCA5 by the Transcription Factor TCP4 Integrates Developmental and Environmental Signals to Promote Hypocotyl Elongation in Arabidopsis. Plant Cell 28, 2117–2130, doi/10.1105/tpc.16.00360 [doi].

Chen, Q., Dai, X., De-Paoli, H., Cheng, Y., Takebayashi, Y., Kasahara, H., Kamiya, Y., and Zhao, Y. (2014). Auxin overproduction in shoots cannot rescue auxin deficiencies in Arabidopsis roots. Plant Cell Physiol. 55, 1072–1079, doi/10.1093/pcp/pcu039 [doi].

Cheng, Y., Dai, X., and Zhao, Y. (2006). Auxin biosynthesis by the YUCCA flavin monooxygenases controls the formation of floral organs and vascular tissues in Arabidopsis. Genes Dev. 20, 1790–1799.

Clough, S.J. and Bent, A.F. (1998). Floral dip: a simplified method for Agrobacterium-mediated transformation of Arabidopsis thaliana. 16, 735–743.

Copeland, N.G., Jenkins, N.A., and Court, D.L. (2001). Recombineering: a powerful new tool for mouse functional genomics. Nat. Rev. Genet. 2, 769–779, doi/10.1038/35093556.

Dahan-Meir, T., Filler-Hayut, S., Melamed-Bessudo, C., Bocobza, S., Czosnek, H., Aharoni, A., and Levy, A.A. (2018). Efficient in planta gene targeting in tomato using geminiviral replicons and the CRISPR/Cas9 system. Plant J. 95, 5–16, doi/10.1111/tpj.13932 [doi].

Ejsmont, R.K., Sarov, M., Winkler, S., Lipinski, K.A., and Tomancak, P. (2009). A toolkit for high-throughput, cross-species gene engineering in Drosophila. Nat. Methods 6, 435–437, doi/10.1038/nmeth.1334.

Fabregas, N., Formosa-Jordan, P., Confraria, A., Siligato, R., Alonso, J.M., Swarup, R., Bennett, M.J., Mahonen, A.P., Cano-Delgado, A.I., and Ibanes, M. (2015). Auxin influx carriers control vascular patterning and xylem differentiation in Arabidopsis thaliana. PLoS Genet. 11, e1005183, doi/10.1371/journal.pgen.1005183 [doi].

Gomez, M.D., Fuster-Almunia, C., Ocana-Cuesta, J., Alonso, J.M., and Perez-Amador, M.A. (2019). RGL2 controls flower development, ovule number and fertility in Arabidopsis. Plant Sci. 281, 82–92, doi/S0168-9452(18)31148-8 [pii].

Grefen, C., Donald, N., Hashimoto, K., Kudla, J., Schumacher, K., and Blatt, M.R. (2010). A ubiquitin-10 promoter-based vector set for fluorescent protein tagging facilitates temporal stability and native protein distribution in transient and stable expression studies. Plant J. 64, 355–365, doi/10.1111/j.1365-313X.2010.04322.x [doi].

Hamilton, C.M. (1997). A binary-BAC system for plant transformation with high-molecular-weight DNA. Gene 200, 107–116, doi/S0378-1119(97)00388-0 [pii].

Han, S.W., Alonso, J.M., and Rojas-Pierce, M. (2015). REGULATOR OF BULB BIOGENESIS1 (RBB1) Is Involved in Vacuole Bulb Formation in Arabidopsis. PLoS One 10, e0125621, doi/10.1371/journal.pone.0125621 [doi].

Hirose, Y., Suda, K., Liu, Y.G., Sato, S., Nakamura, Y., Yokoyama, K., Yamamoto, N., Hanano, S., Takita, E., Sakurai, N., Suzuki, H., Nakamura, Y., Kaneko, T., Yano, K., Tabata, S., and Shibata, D. (2015). The Arabidopsis TAC Position Viewer: a high-resolution map of transformation-competent artificial chromosome (TAC) clones aligned with the Arabidopsis thaliana Columbia-0 genome. Plant J. 83, 1114–1122, doi/10.1111/tpj.12949 [doi].

Isaacs, F.J., Carr, P.A., Wang, H.H., Lajoie, M.J., Sterling, B., Kraal, L., Tolonen, A.C., Gianoulis, T.A., Goodman, D.B., Reppas, N.B., Emig, C.J., Bang, D., Hwang, S.J., Jewett, M.C., Jacobson, J.M., and Church, G.M. (2011). Precise manipulation of chromosomes in vivo enables genome-wide codon replacement. Science 333, 348–353, doi/10.1126/science.1205822 [doi].

Kasahara, H. (2016). Current aspects of auxin biosynthesis in plants. Biosci. Biotechnol. Biochem. 80, 34–42, doi/10.1080/09168451.2015.1086259 [doi].

Kriechbaumer, V., Wang, P., Hawes, C., and Abell, B.M. (2012). Alternative splicing of the auxin biosynthesis gene YUCCA4 determines its subcellular compartmentation. Plant J. 70, 292–302, doi/10.1111/j.1365-313X.2011.04866.x [doi].

Kurihara, D., Mizuta, Y., Sato, Y., and Higashiyama, T. (2015). ClearSee: a rapid optical clearing reagent for whole-plant fluorescence imaging. Development 142, 4168–4179, doi/10.1242/dev.127613 [doi].

Lee, M., Jung, J.H., Han, D.Y., Seo, P.J., Park, W.J., and Park, C.M. (2012). Activation of a flavin monooxygenase gene YUCCA7 enhances drought resistance in Arabidopsis. Planta 235, 923–938, doi/10.1007/s00425-011-1552-3 [doi].

Li, J., Zhang, X., Sun, Y., Zhang, J., Du, W., Guo, X., Li, S., Zhao, Y., and Xia, L. (2018). Efficient allelic replacement in rice by gene editing: A case study of the NRT1.1B gene. J. Integr. Plant. Biol. 60, 536–540, doi/10.1111/jipb.12650 [doi].

Liu, Y.G., Liu, H., Chen, L., Qiu, W., Zhang, Q., Wu, H., Yang, C., Su, J., Wang, Z., Tian, D., and Mei, M. (2002). Development of new transformation-competent artificial chromosome vectors and rice genomic libraries for efficient gene cloning. Gene 282, 247–255.

Mashiguchi, K., Tanaka, K., Sakai, T., Sugawara, S., Kawaide, H., Natsume, M., Hanada, A., Yaeno, T., Shirasu, K., Yao, H., McSteen, P., Zhao, Y., Hayashi, K., Kamiya, Y., and Kasahara, H. (2011). The main auxin biosynthesis pathway in Arabidopsis. Proc. Natl. Acad. Sci. U. S. A. 108, 18512–18517, doi/10.1073/pnas.1108434108.

Peret, B., Swarup, K., Ferguson, A., Seth, M., Yang, Y., Dhondt, S., James, N., Casimiro, I., Perry, P., Syed, A., Yang, H., Reemmer, J., Venison, E., Howells, C., Perez-Amador, M.A., Yun, J., Alonso, J., Beemster, G.T., Laplaze, L., Murphy, A., Bennett, M.J., Nielsen, E., and Swarup, R. (2012a). AUX/LAX genes encode a family of auxin influx transporters that perform distinct functions during Arabidopsis development. Plant Cell 24, 2874–2885, doi/10.1105/tpc.112.097766 [doi].

Peret, B., Swarup, K., Ferguson, A., Seth, M., Yang, Y., Dhondt, S., James, N., Casimiro, I., Perry, P., Syed, A., Yang, H., Reemmer, J., Venison, E., Howells, C., Perez-Amador, M.A., Yun, J., Alonso, J., Beemster, G.T., Laplaze, L., Murphy, A., Bennett, M.J., Nielsen, E., and Swarup, R. (2012b). AUX/LAX genes encode a family of auxin influx transporters that perform distinct functions during Arabidopsis development. Plant Cell 24, 2874–2885, doi/10.1105/tpc.112.097766 [doi].

Pietra, S., Gustavsson, A., Kiefer, C., Kalmbach, L., Horstedt, P., Ikeda, Y., Stepanova, A.N., Alonso, J.M., and Grebe, M. (2013). Arabidopsis SABRE and CLASP interact to stabilize cell division plane orientation and planar polarity. Nat. Commun. 4, 2779, doi/10.1038/ncomms3779 [doi].

Poser, I., Sarov, M., Hutchins, J.R., Heriche, J.K., Toyoda, Y., Pozniakovsky, A., Weigl, D., Nitzsche, A., Hegemann, B., Bird, A.W., Pelletier, L., Kittler, R., Hua, S., Naumann, R., Augsburg, M., Sykora, M.M., Hofemeister, H., Zhang, Y., Nasmyth, K., White, K.P., Dietzel, S., Mechtler, K., Durbin, R., Stewart, A.F., Peters, J.M., Buchholz, F., and Hyman, A.A. (2008). BAC TransgeneOmics: a high-throughput method for exploration of protein function in mammals. Nat. Methods 5, 409–415, doi/10.1038/nmeth.1199.

Ruzicka, K., Ljung, K., Vanneste, S., Podhorska, R., Beeckman, T., Friml, J., and Benkova, E. (2007). Ethylene Regulates Root Growth through Effects on Auxin Biosynthesis and Transport-Dependent Auxin Distribution. Plant Cell 19, 2197–2212.

Sabatini, S., Beis, D., Wolkenfelt, H., Murfett, J., Guilfoyle, T., Malamy, J., Benfey, P., Leyser, O., Bechtold, N., Weisbeek, P., and Scheres, B. (1999). An auxin-dependent distal organizer of pattern and polarity in the Arabidopsis root. Cell 99, 463–472.

Sandvang, D. (1999). Novel streptomycin and spectinomycin resistance gene as a gene cassette within a class 1 integron isolated from Escherichia coli. Antimicrob. Agents Chemother. 43, 3036–3038.

Sarov, M., Schneider, S., Pozniakovski, A., Roguev, A., Ernst, S., Zhang, Y., Hyman, A.A., and Stewart, A.F. (2006). A recombineering pipeline for functional genomics applied to Caenorhabditis elegans. Nat. Methods 3, 839–844, doi/10.1038/nmeth933.

Sarov, M., Murray, J.I., Schanze, K., Pozniakovski, A., Niu, W., Angermann, K., Hasse, S., Rupprecht, M., Vinis, E., Tinney, M., Preston, E., Zinke, A., Enst, S., Teichgraber, T., Janette, J., Reis, K., Janosch, S., Schloissnig, S., Ejsmont, R.K., Slightam, C., Xu, X., Kim, S.K., Reinke, V., Stewart, A.F., Snyder, M., Waterston, R.H., and Hyman, A.A. (2012). A genome-scale resource for in vivo tag-based protein function exploration in C. elegans. Cell 150, 855–866, doi/10.1016/j.cell.2012.08.001; 10.1016/j.cell.2012.08.001.

Sarov, M., Barz, C., Jambor, H., Hein, M.Y., Schmied, C., Suchold, D., Stender, B., Janosch, S., K, J.V.V., Krishnan, R.T., Krishnamoorthy, A., Ferreira, I.R., Ejsmont, R.K., Finkl, K., Hasse, S., Kampfer, P., Plewka, N., Vinis, E., Schloissnig, S., Knust, E., Hartenstein, V., Mann, M., Ramaswami, M., VijayRaghavan, K., Tomancak, P., and Schnorrer, F. (2016). A genome-wide resource for the analysis of protein localisation in Drosophila. Elife 5, e12068, doi/10.7554/eLife.12068 [doi].

Schlake, T. and Bode, J. (1994). Use of mutated FLP recognition target (FRT) sites for the exchange of expression cassettes at defined chromosomal loci. Biochemistry 33, 12746–12751.

Smyth, D.R., Bowman, J.L., and Meyerowitz, E.M. (1990). Early flower development in Arabidopsis. Plant Cell 2, 755–767, doi/10.1105/tpc.2.8.755 [doi].

Soyars, C.L., Peterson, B.A., Burr, C.A., and Nimchuk, Z.L. (2018). Cutting Edge Genetics: CRISPR/Cas9 Editing of Plant Genomes. Plant Cell Physiol. 59, 1608–1620, doi/10.1093/pcp/pcy079 [doi].

Stepanova, A.N., Robertson-Hoyt, J., Yun, J., Benavente, L.M., Xie, D., Dolezal, K., Schlereth, A., Jurgens, G., and Alonso, J.M. (2008). TAA1-mediated auxin biosynthesis is essential for hormone crosstalk and plant development Cell 133, 177–191.

Stepanova, A.N., Hoyt, J.M., Hamilton, A.A., and Alonso, J.M. (2005). A Link between ethylene and auxin uncovered by the characterization of two root-specific ethylene-insensitive mutants in Arabidopsis. Plant Cell 17, 2230–2242.

Stepanova, A.N., Yun, J., Likhacheva, A.V., and Alonso, J.M. (2007). Multilevel interactions between ethylene and auxin in Arabidopsis roots. Plant Cell 19, 2169–2185.

Stepanova, A.N., Yun, J., Robles, L.M., Novak, O., He, W., Guo, H., Ljung, K., and Alonso, J.M. (2011). The Arabidopsis YUCCA1 Flavin Monooxygenase Functions in the Indole-3-Pyruvic Acid Branch of Auxin Biosynthesis. Plant Cell 23, 3961–3973, doi/10.1105/tpc.111.088047.

Sugawara, S., Hishiyama, S., Jikumaru, Y., Hanada, A., Nishimura, T., Koshiba, T., Zhao, Y., Kamiya, Y., and Kasahara, H. (2009). Biochemical analyses of indole-3-acetaldoxime-dependent auxin biosynthesis in Arabidopsis. Proc. Natl. Acad. Sci. U. S. A., doi/10.1073/pnas.0811226106.

Swarup, R., Perry, P., Hagenbeek, D., Van Der Straeten, D., Beemster, G.T., Sandberg, G., Bhalerao, R., Ljung, K., and Bennett, M.J. (2007). Ethylene upregulates auxin biosynthesis in Arabidopsis seedlings to enhance inhibition of root cell elongation. Plant Cell 19, 2186–2196.

Tam, Y.Y. and Normanly, J. (1998). Determination of indole-3-pyruvic acid levels in Arabidopsis thaliana by gas chromatography-selected ion monitoring-mass spectrometry. J. Chromatogr. A 800, 101–108, doi/S0021-9673(97)01051-0 [pii].

Tao, Y., Ferrer, J.L., Ljung, K., Pojer, F., Hong, F., Long, J.A., Li, L., Moreno, J.E., Bowman, M.E., Ivans, L.J., Cheng, Y., Lim, J., Zhao, Y., Ballare, C.L., Sandberg, G., Noel, J.P., and Chory, J. (2008). Rapid synthesis of auxin via a new tryptophan-dependent pathway is required for shade avoidance in plants. Cell 133, 164–176, doi/10.1016/j.cell.2008.01.049.

Tian, G.W., Mohanty, A., Chary, S.N., Li, S., Paap, B., Drakakaki, G., Kopec, C.D., Li, J., Ehrhardt, D., Jackson, D., Rhee, S.Y., Raikhel, N.V., and Citovsky, V. (2004). High-throughput fluorescent tagging of full-length Arabidopsis gene products in planta. Plant Physiol. 135, 25–38.

Tiwari, S.B., Wang, X.J., Hagen, G., and Guilfoyle, T.J. (2001). AUX/IAA proteins are active repressors, and their stability and activity are modulated by auxin. Plant Cell 13, 2809–2822, doi/10.1105/tpc.010289 [doi].

Turan, S., Zehe, C., Kuehle, J., Qiao, J., and Bode, J. (2013). Recombinase-mediated cassette exchange (RMCE) - a rapidly-expanding toolbox for targeted genomic modifications. Gene 515, 1–27, doi/10.1016/j.gene.2012.11.016 [doi].

Tursun, B., Cochella, L., Carrera, I., and Hobert, O. (2009). A toolkit and robust pipeline for the generation of fosmid-based reporter genes in C. elegans. PLoS One 4, e4625, doi/10.1371/journal.pone.0004625.

Vanneste, S. and Friml, J. (2009). Auxin: a trigger for change in plant development. Cell 136, 1005–1016, doi/10.1016/j.cell.2009.03.001.

Vaseva, I.I., Qudeimat, E., Potuschak, T., Du, Y., Genschik, P., Vandenbussche, F., and Van Der Straeten, D. (2018). The plant hormone ethylene restricts Arabidopsis growth via the epidermis. Proc. Natl. Acad. Sci. U. S. A. 115, E4130–E4139, doi/10.1073/pnas.1717649115 [doi].

Venken, K.J., He, Y., Hoskins, R.A., and Bellen, H.J. (2006). P[acman]: a BAC transgenic platform for targeted insertion of large DNA fragments in D. melanogaster. Science 314, 1747–1751, doi/10.1126/science.1134426.

Venken, K.J., Kasprowicz, J., Kuenen, S., Yan, J., Hassan, B.A., and Verstreken, P. (2008). Recombineering-mediated tagging of Drosophila genomic constructs for in vivo localization and acute protein inactivation. Nucleic Acids Res. 36, e114, doi/10.1093/nar/gkn486.

Venken, K.J., Carlson, J.W., Schulze, K.L., Pan, H., He, Y., Spokony, R., Wan, K.H., Koriabine, M., de Jong, P.J., White, K.P., Bellen, H.J., and Hoskins, R.A. (2009). Versatile P[acman] BAC libraries for transgenesis studies in Drosophila melanogaster. Nat. Methods 6, 431–434, doi/10.1038/nmeth.1331.

Villarino, G.H., Hu, Q., Manrique, S., Flores-Vergara, M., Sehra, B., Robles, L., Brumos, J., Stepanova, A.N., Colombo, L., Sundberg, E., Heber, S., and Franks, R.G. (2016). Transcriptomic Signature of the SHATTERPROOF2 Expression Domain Reveals the Meristematic Nature of Arabidopsis Gynoecial Medial Domain. Plant Physiol. 171, 42–61, doi/10.1104/pp.15.01845 [doi].

Wang, Y. and Jiao, Y. (2018). Auxin and above-ground meristems. J. Exp. Bot. 69, 147–154, doi/10.1093/jxb/erx299 [doi].

Warming, S., Costantino, N., Court, D.L., Jenkins, N.A., and Copeland, N.G. (2005). Simple and highly efficient BAC recombineering using galK selection. Nucleic Acids Res. 33, e36.

Worden, N., Wilkop, T.E., Esteve, V.E., Jeannotte, R., Lathe, R., Vernhettes, S., Weimer, B., Hicks, G., Alonso, J., Labavitch, J., Persson, S., Ehrhardt, D., and Drakakaki, G. (2015). CESA TRAFFICKING INHIBITOR inhibits cellulose deposition and interferes with the trafficking of cellulose synthase complexes and their associated proteins KORRIGAN1 and POM2/CELLULOSE SYNTHASE INTERACTIVE PROTEIN1. Plant Physiol. 167, 381–393, doi/10.1104/pp.114.249003 [doi].

Xu, Y., Prunet, N., Gan, E.S., Wang, Y., Stewart, D., Wellmer, F., Huang, J., Yamaguchi, N., Tatsumi, Y., Kojima, M., Kiba, T., Sakakibara, H., Jack, T.P., Meyerowitz, E.M., and Ito, T. (2018). SUPERMAN regulates floral whorl boundaries through control of auxin biosynthesis. EMBO J. 37, 10.15252/embj.201797499. Epub 2018 May 15, doi/e97499 [pii].

Yamada, M., Greenham, K., Prigge, M.J., Jensen, P.J., and Estelle, M. (2009). The TRANSPORT INHIBITOR RESPONSE2 gene is required for auxin synthesis and diverse aspects of plant development. Plant Physiol. 151, 168–179, doi/10.1104/pp.109.138859.

Yanagisawa, M., Alonso, J.M., and Szymanski, D.B. (2018). Microtubule-Dependent Confinement of a Cell Signaling and Actin Polymerization Control Module Regulates Polarized Cell Growth. Curr. Biol. 28, 2459–2466.e4, doi/S0960-9822(18)30712-7 [pii].

Yu, D., Ellis, H.M., Lee, E.C., Jenkins, N.A., Copeland, N.G., and Court, D.L. (2000). An efficient recombination system for chromosome engineering in Escherichia coli. Proc. Natl. Acad. Sci. U. S. A. 97, 5978–5983.

Yu, Q.H., Wang, B., Li, N., Tang, Y., Yang, S., Yang, T., Xu, J., Guo, C., Yan, P., Wang, Q., and Asmutola, P. (2017). CRISPR/Cas9-induced Targeted Mutagenesis and Gene Replacement to Generate Long-shelf Life Tomato Lines. Sci. Rep. 7, 11874–017-12262-1, doi/10.1038/s41598-017-12262-1 [doi].

Yuan, Q., Liang, F., Hsiao, J., Zismann, V., Benito, M.I., Quackenbush, J., Wing, R., and Buell, R. (2000). Anchoring of rice BAC clones to the rice genetic map in silico. Nucleic Acids Res. 28, 3636–3641, doi/10.1093/nar/28.18.3636 [doi].

Zhang, J., Chen, L.L., Sun, S., Kudrna, D., Copetti, D., Li, W., Mu, T., Jiao, W.B., Xing, F., Lee, S., Talag, J., Song, J.M., Du, B., Xie, W., Luo, M., Maldonado, C.E., Goicoechea, J.L., Xiong, L., Wu, C., Xing, Y., Zhou, D.X., Yu, S., Zhao, Y., Wang, G., Yu, Y., Luo, Y., Hurtado, B.E., Danowitz, A., Wing, R.A., and Zhang, Q. (2016). Building two indica rice reference genomes with PacBio long-read and Illumina paired-end sequencing data. Sci. Data 3, 160076, doi/10.1038/sdata.2016.76 [doi].

Zhang, T.Q., Xu, Z.G., Shang, G.D., and Wang, J.W. (2019). A Single-Cell RNA Sequencing Profiles the Developmental Landscape of Arabidopsis Root. Mol. Plant. 12, 648–660, doi/S1674-2052(19)30133-9 [pii].

Zhao, Y. (2018). Essential Roles of Local Auxin Biosynthesis in Plant Development and in Adaptation to Environmental Changes. Annu. Rev. Plant. Biol. 69, 417–435, doi/10.1146/annurev-arplant-042817-040226 [doi].

Zhou, R., Benavente, L.M., Stepanova, A.N., and Alonso, J.M. (2011). A recombineering-based gene tagging system for Arabidopsis. Plant J. 66, 712–723, doi/10.1111/j.1365-313X.2011.04524.x; 10.1111/j.1365-313X.2011.04524.x.

